# A quantitative review of abundance-based species distribution models

**DOI:** 10.1101/2021.05.25.445591

**Authors:** Conor Waldock, Rick D. Stuart-Smith, Camille Albouy, William W. L. Cheung, Graham J. Edgar, David Mouillot, Jerry Tjiputra, Loïc Pellissier

## Abstract

The contributions of species to ecosystem functions or services depend not only on their presence in a given community, but also on their local abundance. Progress in predictive spatial modelling has largely focused on species occurrence, rather than abundance. As such, limited guidance exists on the most reliable methods to explain and predict spatial variation in abundance. We analysed the performance of 68 abundance-based species distribution models fitted to 800,000 standardised abundance records for more than 800 terrestrial bird and reef fish species. We found high heterogeneity in performance of abundance-based models. While many models performed poorly, a subset of models consistently reconstructed range-wide abundance patterns. The best predictions were obtained using random forests for frequently encountered and abundant species, and for predictions within the same environmental domain as model calibration. Extending predictions of species abundance outside of the environmental conditions used in model training generated poor predictions. Thus, interpolation of abundances between observations can help improve understanding of spatial abundance patterns, but extrapolated predictions of abundance, e.g. under climate change, have a much greater uncertainty. Our synthesis provides a roadmap for modelling abundance patterns, a key property of species’ distributions that underpins theoretical and applied questions in ecology and conservation.

## Introduction

Environmental change alters the occurrence and local abundance patterns of species (Hastings et al. 2020, Román-Palacios and Wiens 2020, Lenoir et al. 2020, Antão et al. 2020b). Modelling species’ occurrence has helped predict the distribution and erosion of biodiversity under unprecedented rates of environmental change (Pereira et al. 2013, Kissling et al. 2018, Jetz et al. 2019). Species occurrence models, however, provide limited opportunities to understand local abundance changes that accompany species distribution shifts (Lenoir and Svenning 2013, Bates et al. 2015, Hastings et al. 2020). Species present in high numbers at only a few sites can make large contributions to ecological processes but a focus on occurrence would overlook these species (Table 1: (Stuart-Smith et al. 2013, Williams et al. 2014, Winfree et al. 2015, Johnston et al. 2015, Genung et al. 2020)). Abundance trends can also act as an early warning signal of population collapse (Clements et al. 2017, Ceballos et al. 2020) but occurrence patterns may not change until after local population depletion (Hastings et al. 2020). To better inform spatial conservation planning, we must better monitor and predict species abundance (Margules and Pressey 2000, Pauly and Froese 2010, Mi et al. 2017); however, abundance-based species distribution models remain under-developed relative to occurrence-based models.

**Table 1.** Role of species’ abundance information in applied ecology and conservation.

| Research topic | Benefit of abundance information | Application |
| --- | --- | --- |
| Monitoring biodiversity change | Population and patch extinction risk is better predicted by patch abundance rather than occupancy alone. | Schulz et al. (2020) show abundance in the previous year to be a strong predictor of Glanville fritillary (*Melitaea cinxia*) butterfly patch occupancy, such that local abundance rather than average abundance determines local extinction risks. |
|  | If using a fixed focal area for surveys, species' environmental response curves are better quantified using abundance, which provides more information than presence-absence. | Becker et al. (2019) modelled the influx of cetacean individuals to the California current system, using generalised additive models, during a heatwave event of 2014. |
|  | Quantitative changes in abundance within a species range are more informative than occurrence shifts (i.e., intermediate stages in range shifts, no change in range extent). | Fei et al. (2017) found that shifts in the spatial distribution of species' abundance for tree species in the United States from 1980s to 2010s, was mostly due to sub-populations increasing in density from low initial abundance. |
|  | Abundance is more sensitive at detecting impacts on species' distributions than occurrence. | Maxwell et al. (2019) synthesised 698 studies responses to extreme weather events and showed that abundance declines occurred in 100 cases, but local extinction occurred in only 31 cases. Ricart et al. (2018) show that habitat forming *Codium vermilara* algae in the north west Mediterranean has declined by 95% in terms of abundance but only 45% in terms of site occupancy. |
|  | Trends in abundance and species richness can be disconnected. | Antão et al. (2020a) found contrasting patterns in assemblage abundance and species richness in Finnish moth assemblages over 19 years, with abundance declining despite species richness increasing. |
| Ecosystem function and services | Individuals contribute to ecosystem services rather than species. | Winfree et al. (2015) found that, in real-world ecosystems, crop pollination was driven by abundance fluctuations of dominant bee species whereas species richness was driven by rare species that contributed little to ecosystem function. |
|  | Interaction strengths depend on the abundance of interacting species. | Matías et al. (2019) document how pathogen abundance determines Cork oak (*Quercus suber*) mortality rates across the species' distribution. More generally, Vázquez et al. (2007) show that asymmetry in interaction strength between hosts and consumers is correlated with abundance, so that rarer species are more negatively affected by abundant partners but pairs of interacting abundant species exhibit reciprocally strong effects. |
|  | Geographic differences in patterns in evenness in abundance exist, such that the contributions of individuals and species to assemblage functional diversity varies at a macroecological scale. | Stuart-Smith et al. (2013) show that community evenness is higher in temperate reef fish assemblages, compared to tropical assemblages. This difference in assemblage evenness suggest that each fish species contribution to reef ecosystem functioning is higher in temperate than tropical regions. |
|  | Productivity depends on number of individuals in an area, which can map differently to the area suitable for occupancy. | Kallasvuori et al. (2017) demonstrate that the most productive areas, with most individuals, only occupy a small area of the total suitable region for fish stocks in the Baltic Sea. |
| Management of biodiversity | Management goals are often to maintain abundance (biomass) of individuals rather than just presence | Hutchings and Reynolds (2004) show breeding population sizes of economically valuable fishes have declined by 83%, undermining profitable fisheries, even though small populations still persist. |
|  | Extinction risk is often established based on population abundance change, which can be spatially variable | Sherley et al. (2020) use 40 years of count data of African penguin (*Spheniscus demersus*) and model spatially dependent abundance change through time to identify regions in the geographic range at high risk of extinction. The overall decline in abundance was 65% since 1989, indicating that the threshold for the IUCN 'Endangered' Red List category had been crossed. |
|  | Spatial mapping of abundance for prioritization of area of conservation | Flores et al. (2018) show how valley areas are important for maintaining high populations of Guanaco (*Lama guanicoe*) in central Tierra del Fuego, and that spatial heterogeneity of abundance is greater in the breeding than non-breeding season. |
|  | Invader impact curves suggest impacts are threshold dependent. | Yokomizo et al. (2009) simulations indicate that impacts of invasive species depend on density, and that density-impact curve must be correctly identified to prevent overinvestment in management with little reduction in impact, particularly for species whose impact is only realised at high densities. |

As in occurrence-based models, modelling abundance according to abiotic environmental conditions depends on assumptions of niche theory (Maguire, 1973, Holt 2009). Critically, environmental conditions are assumed to affect demographic processes which in turn drive population dynamics (Maguire, 1973, Brown et al. 1995, Holt 2009, Pearce-Higgins et al. 2015, Betts et al. 2019). For a given species, spatial abundance variation is a consequence of these links coupled with natural environmental gradients (Holt 2009). If this theory is accurate, predictions of local abundance from environmental factors should be possible (Maguire, 1973, Martínez-Meyer et al. 2013, Waldock et al. 2019).

Yet, abundance does not appear to always be strongly constrained by theoretical niche properties in empirical data (Yañez-Arenas et al. 2014a, Dallas et al. 2017, Osorio-olvera et al. 2019, Santini et al. 2019, Dallas and Santini 2020, Holt 2020, Sporbert et al. 2020). For example, Allee effects, non-equilibrium population states, demographic stochasticity, and environmental variability act to weaken the link between environmental conditions and local abundance (Osorio-olvera et al. 2019, Dallas and Santini 2020, Holt 2020). If these factors dominate over macro-environmental constraints on abundance, then abundance will be poorly predicted using a species distribution modelling approach. At present, the expected predictive power when modelling abundance in relation to environmental conditions is poorly understood and not quantitatively reviewed over large datasets and a varied set of modelling frameworks.

Recent decades of statistical algorithm development provide an opportunity to evaluate the performance of abundance-based species distribution models. Current abundance model evaluations examine only a limited set of statistical frameworks and the best options may be overlooked (Pearce and Ferrier 2001, Potts and Elith 2006, Oppel et al. 2012, Bahn and McGill 2013). For example, if abundance is determined by non-linear and complex interactions of environmental factors, then machine-learning algorithms may be most appropriate (Merow et al. 2014, Damaris et al. 2016). In contrast, simpler models may be favoured if a species’ environmental responses closely follows simple unimodal functions (Austin 2002, Ready et al. 2010, Boucher-Lalonde et al. 2012, Waldock et al. 2019). Simpler models are also expected to perform better when extrapolated to new environmental conditions (Merow et al. 2014, Brun et al. 2019).

The scarcity of abundance data across entire species ranges has likely also contributed to poor model development (i.e., a Prestonian shortfall; (Pauly and Froese 2010, Cardoso et al. 2011, Hortal et al. 2015)). However, the technological expansion in citizen-science has generated a rapidly increasing quantity of species’ abundance records (Dickinson et al. 2010, Edgar and Stuart-Smith 2014), which combined with many national and regional biomonitoring surveys could allow the large-scale application of abundance-based species distribution models (Margules and Pressey 2000, Kissling et al. 2018, Callaghan et al. 2021).

Species distribution model performance is often associated with species and data characteristics. Establishing how and why model performance varies for different species is critical for conservation and management applications, particularly with respect to commonness. Common species, in terms of local and regional abundance, often contribute most to ecosystem functioning (Genung et al. 2020). Low abundance and range-restricted species may be prioritised for conservation, having higher extinction risk (Purvis et al. 2000, Ceballos et al. 2020) and potentially playing unique roles in ecosystems (Violle et al. 2017). Species distribution models generally perform better for species with smaller ranges, lower endemicity and non-migratory behaviour, in addition, the number of observations positively affects performance (McPherson and Jetz 2007, Newbold et al. 2009, Chefaoui et al. 2011, Thuiller et al. 2019). The influence of species characteristics on abundance model performance is less well established. Furthermore, in novel environmental conditions the species characteristics associated with extrapolating abundance predictions are important to identify.

Effects of species characteristics in abundance-based models may differ from occurrence-based models. Differences could arise because species abundance is jointly determined by fundamental niche axes in addition to dynamic population properties (Peterson et al. 2011), such as the strength of negative density dependence, intrinsic population growth rates and population cycles (Chisholm and Muller-Landau 2011, Yañez-Arenas et al. 2014b, Chu et al. 2016, Bowler et al. 2017, Yenni et al. 2017, Hallett et al. 2018). Fundamental niche limits are expected to play a small role in controlling abundance of wide-ranging species, because these species have their abundance controlled by a milieu of demographic factors that may each have different response functions (Hallett et al. 2018), perhaps leading to lower performance. In contrast, rare (low mean abundance) species that have narrow niches are theoretically expected to exhibit more stable populations and could therefore exhibit more predictable abundances (Yenni et al. 2017).

Data characteristics, such as the amount of observations, are another element that could affect the success of species distribution model performance (Wisz et al. 2008, Yañez-Arenas et al. 2014b). More samples generally improve species distribution model performance by being less geographically and environmentally biased (Wisz et al. 2008), and should similarly improve abundance model performance (Yañez-Arenas et al. 2014b). Yet, these effects have not been tested.

Here, we aim to provide practical guidance on applying statistical approaches to predict species’ abundance, and identify factors most affecting predictive performance. We compare 68 abundance-based species distribution models fitted for two standardised abundance datasets containing more than 800 marine and terrestrial vertebrate species and over 800,000 abundance observations. We test model interpolative (within-sample) and extrapolative (out-of-sample) performance. We ask how statistical framework and model complexity, and species’ and data characteristics, affect metrics of model accuracy, discrimination, and precision. We show that abundance-based species distribution models have great potential – additional to occurrence-based models – to generate insights in spatial ecology and biogeography, and to improve systematic conservation planning outcomes.

## Materials and methods

### Spatial abundance data

We obtained standardised estimates of species abundance across large regions for birds and shallow-water reef fishes from the Breeding Bird Survey of the USA (BBS) and Reef Life Survey (RLS) respectively (for detailed sampling schemes see (Pardieck et al. 2019) for birds and (Edgar and Stuart-Smith 2014) for fishes). For birds, abundance data comprise of 3-minute counts of individuals sighted and heard within a 400m stop radius along a transect of 50 stops. We summed bird species’ abundance across 50-stops within a sampled year, and mean-averaged abundances for a given species in a repeated site across the years 2014-2018. We aggregated abundances across years to better generalise our results to the structure of most abundance datasets, whereby yearly values across broad geographic regions are unlikely to be available (see Figure S25 for exploration of this assumption). We filtered out all samples that did not meet BBS established weather, date, time, route completion, randomised sampling, and sampling protocol criteria (i.e., using BBS data with a run type of 1). For fishes, abundance data are counts of individuals sighted along 50m long underwater transects (summed across 2 x 5m wide blocks either side of the transect line). We mean-averaged RLS abundance estimates across multiple transects within sites and we defined sites as sets of transects <200m apart (Edgar and Stuart-Smith 2014, Cresswell et al. 2017). We filtered sites geographically between 3°S to 50°S and 110°E to 165°E to select for Australian and Indo-Pacific survey locations where sampling effort was most intensive and comprehensive in the RLS dataset. For both BBS and RLS datasets, we removed species without full scientific names and fewer than 50 abundance records. We required species absences for two-stage models and abundance-absence models. We generated absences for each species by taking observations where species were present and finding all observations within a 1000 km buffer where species were not present. A lack of observed presence is not necessarily a ‘true absence’, but instead suggests species were undetectable with a reasonable sampling effort (Guillera-Arroita 2017). We analysed a total of 264,474 observations of 385 species in 3,890 sites for birds in the BBS dataset, and 567,669 observations of 495 species in 2,137 sites for reef fishes in the RLS dataset.

### Covariates

We matched site locations to gridded environmental variables representing climate, biogeochemistry, land-use, depth, habitat area, and human populations, retaining only variables with expected *a priori* relationships with abundance (see Table S1 for details). Because of the high number of similar climate-related variables, and to avoid multicollinearity in these, we first applied robust-PCA using package *pcaMethods* (version 1.76.0; Stacklies et al. 2007) which is shown to be a good approach to reduce multicollinearity in SDMs (Cruz-Cárdenas et al. 2014, De Marco and Nóbrega 2018, Osorio-Olvera et al. 2020). Furthermore, we focused on predictive power to ensure our results were more robust to potential multicollinearity. We ran a separate robust-PCA on 19 variables characterising climates across the bird survey locations (bio1-bio19), and on 15 variables characterising climatic and biogeochemical properties across the fish survey locations (mean, minimum and maximum of pH, salinity, chl-a, net primary productivity, degree heating weeks; indicated in Table S1). For each dataset, we retained 3 principal components, explaining 87.8% and 77.8% variation respectively, and used these principal component scores as predictor variables to summarise the dominant climate and biogeochemical regimes of the data in each set of models (3 PCA variables for birds and 3 PCA variables for fishes; Figure S1 and S2). We retained the PCA axes which explained >5% variation in the PCA-covariate set, which resulted in 3 axes summarising the climatological variation for each dataset. In addition to the climatological variables, we also included additional environmental variables as predictors in our model that we expected to act independently. All non-PCA variables were mean-centred, normalised to a variance of 1, and transformed according to Table S1 before modelling.

### Analytical design

We analysed a large diversity of species abundance models that spanned a gradient in model complexity and different formulations of abundance data. Further, we assessed model performance for interpolation and extrapolation cross-validation scenarios (Figure 1). Given that data requirements are a major challenge in fitting species abundance models, we chose species-level statistical models that were suitable for our goal of comparing predictive performance (i.e., not mechanistic, hierarchical or multispecies/joint/multivariate SDMs). In total, we fitted and evaluated 68 types of species abundance model (24 model frameworks by 3 response variable (abundance) formulations, less 4 models of zero-inflation that are not valid for abundance-only models = 68 models; see Table S2 for full model list). Combining models and cross-validations for 1,547 species led to 59,840 models to evaluate.

**Figure 1.**
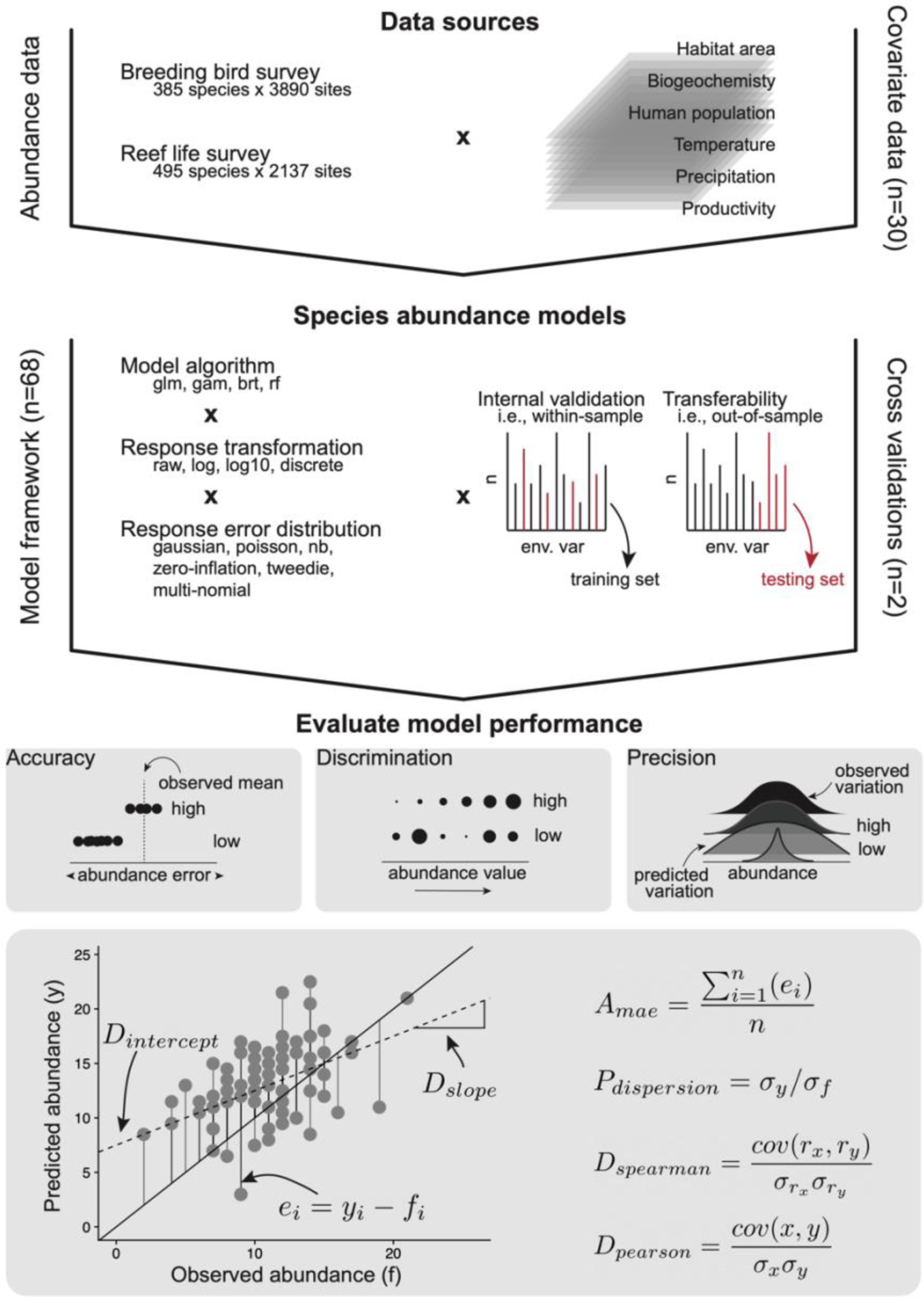
Overview of analysis from data sources to model performance evaluations. Model evaluation metrics for accuracy, discrimination and precision are presented.

**Table 2.** Summary of evaluation metrics of model performance for most discriminatory models comparing within- and out-of-sample cross validations, median and interquartile range (IQR) for all species within datasets are presented. A_mae_ has the proportional error of estimated mean abundance compared to observed mean abundance having a target value of 1. D_pearson_ and D_spearman_ are correlation coefficients having a target value of 1. D_intercept_ is the number of individuals predicted from a linear regression between observed and predicted at 0 observed individuals. D_slope_ is the slope of this regression having a target value of 1. P_dispersion_ is a dimensionless ratio of the standard deviation of predicted abundance over the standard deviation of observed abundance having a target value of 1.

|  | metric | within-sample |  |  |  |  | out-of-sample |  |  |  |  |
| --- | --- | --- | --- | --- | --- | --- | --- | --- | --- | --- | --- |
|  |  | Q0.05 | Q0.25 | median | Q0.75 | Q0.95 | Q0.05 | Q0.25 | median | Q0.75 | Q0.95 |
| breeding<br>bird<br>survey | $A_{mae}$ | 0.43 | 0.54 | 0.62 | 0.70 | 0.97 | 0.47 | 0.65 | 0.78 | 0.92 | 1.88 |
| | $D_{intercept}$ | 0.68 | 1.44 | 2.23 | 3.40 | 9.95 | 0.00 | 0.40 | 1.57 | 4.42 | 19.38 |
| | $D_{slope}$ | 0.02 | 0.15 | 0.25 | 0.36 | 0.68 | 0.00 | 0.05 | 0.10 | 0.19 | 0.45 |
| | $D_{pearson}$ | 0.15 | 0.37 | 0.49 | 0.61 | 0.74 | 0.09 | 0.23 | 0.34 | 0.46 | 0.65 |
| | $D_{spearman}$ | 0.14 | 0.36 | 0.48 | 0.61 | 0.72 | 0.10 | 0.24 | 0.34 | 0.46 | 0.62 |
| | $P_{dispersion}$ | 0.12 | 0.34 | 0.51 | 0.68 | 1.27 | 0.04 | 0.18 | 0.32 | 0.55 | 1.25 |
| reef life<br>survey | $A_{mae}$ | 0.46 | 0.59 | 0.69 | 0.84 | 1.34 | 0.45 | 0.66 | 0.83 | 0.97 | 1.43 |
| | $D_{intercept}$ | 0.25 | 0.90 | 1.68 | 5.75 | 93.70 | -0.01 | 0.42 | 1.50 | 4.68 | 62.63 |
| | $D_{slope}$ | 0.01 | 0.10 | 0.20 | 0.38 | 0.99 | 0.00 | 0.02 | 0.06 | 0.16 | 0.49 |
| | $D_{pearson}$ | 0.15 | 0.33 | 0.48 | 0.63 | 0.84 | 0.05 | 0.21 | 0.36 | 0.50 | 0.74 |
| | $D_{spearman}$ | 0.17 | 0.31 | 0.43 | 0.56 | 0.72 | 0.04 | 0.22 | 0.34 | 0.47 | 0.67 |
| | $P_{dispersion}$ | 0.05 | 0.25 | 0.44 | 0.72 | 1.67 | 0.00 | 0.07 | 0.20 | 0.41 | 1.25 |

Our full species abundance model set comprises different statistical algorithms, response transformations, error distributions, and formulations of abundance data. We used 24 model variants from common statistical distributions and transformations for abundance data that were available within statistical software packages in R (e.g., Poisson, negative binomial, zero-inflated, tweedie, multi-nominal, log10-gaussian, log-gaussian; Table S2). We chose statistical treatments of abundance data that are common in the literature and valid to the error distribution of abundance. We fitted these 24 model variants using four statistical model fitting procedures: generalised linear models (GLM), generalised additive models (GAM; Wood 2011), gradient boosting machine (GBM; Friedman 2001), and random forests (RF; Breiman 2001). This model set varied in complexity of the relationship between abundance and environmental variables (linear to highly-complex) and the behaviour of interactions within the models (none to many; Merow et al. 2014). For GLMs and GAMs we used a range of error distributions rather than determining *a priori* the most appropriate error distribution for each species. This follows previous species abundance model comparisons (Potts and Elith 2006), which assumed that incorrect model specification leads to poor predictive ability, and we focussed our comparison of model performance on predictive ability (which also provided a standardised assessment criteria across statistical algorithms). For all models, we included the same initial set of predictor variables, although each model framework had a different underlying variable selection procedure that identified independent sets of final predictors. The full model fitting procedure, algorithm parameters, and justification for each modelling approach and software used are provided in Appendix 2.

In addition to model variants, we used three formulations of response data: abundance-when-present (for 20 model variants, less 4 zero-inflated models), abundance-absence (for 24 model variants), and an indirect two-stage modelling approach (for 24 model variants). For abundance-when-present models we removed all absences. Abundance-absence models were analogous to classic presence-absence data in species distribution models, but using abundance estimates instead of presences. In abundance-absence models, we standardised prevalence (the number of absences compared to presences) across species, which can influence the estimation of response curves from data characteristics alone when there are many more absences than presences (Meynard et al. 2019). To do so, we bootstrap-subsampled the number of absences to be twice the number of presences, repeating this procedure 10 times and averaging abundance predictions across bootstraps. Finally, our indirect two-stage modelling approach first modelled habitat suitability as a traditional SDM by converting abundance-absences into presence-absences. Next, we used the habitat suitability predictions from this model as a single covariate to predict abundance. Note, this is not a hurdle approach, but instead tests the assumption that habitat suitability correlates to, and predicts, local abundance (Vanderwal et al. 2009). Details for fitting SDMs to produce occupancy predictions are provided in Appendix 2.

### Model evaluation: accuracy, discrimination, and precision

We evaluated the consistency between predicted and observed abundance using metrics of: i) accuracy, ii) discrimination and iii) precision (see Figure1 for equations; (Norberg et al. 2019)). Accuracy is the degree of proximity to a known truth, measured here using mean absolute error between observed and predicted abundance, divided by the mean observed abundance for a species (A_mae_). Discrimination measure how well model predictions discern low values from high values of observed abundance, e.g., in the correct overall ordering of abundances. This is a continuous analogue of occurrence SDMs discerning between present and absent. We measured discrimination using both Spearman’s rank correlation (D_spearman_) and Pearson’s correlation (D_pearson_) between predicted and observed abundance. In addition, we estimated the slope and intercept of a linear model between predicted and observed abundance (D_slope_, D_intercept_). Precision measures the information content in the predictions as the variation in predicted abundance relative to the variation in the observed abundances. Precision differs from accuracy because estimates can be precise with high information content even if overall predictions were biased. Here we measured precision as the standard deviation of the predicted abundances (Norberg et al. 2019). However, we compared this value to a reasonable expectation of precision because each species has a different range of abundance values. Therefore, we estimated the predicted precision divided by the expected variation in abundance and call this property P_dispersion_.

Accuracy, discrimination, and precision capture different facets of model performance and so could be considered together or separately depending on the purposes of the modelling exercise. For example, a model can predict mean abundance of a species well (high accuracy) but poorly discriminate between high and low abundances (low discrimination). We focused our results mostly on discrimination because identifying changes in spatial and temporal variation in abundance, a goal of conservation and wildlife management, depends on good discrimination of abundance values between sites or time-points. Further, accuracy and precision may depend on the quality of sampling, but inaccurate sampling may still provide reasonable estimates of spatial and temporal differences in abundance. We identified an ‘optimal model’ based on the most discriminatory model for each species. To do so, we rescaled the four discrimination metrics between 0-1, averaged the score across the scaled metrics, and identified the model with the highest average score per species – we report this as the ‘optimal model’ throughout.

Note that we avoid confounding performance in predicting presence-absences from performance in predicting abundance by only evaluating predictions for species abundances when present (i.e., we exclude any abundance values predicted in sites where species are absent in the observed data). Many reviews exist identifying the best occupancy based frameworks for predicting presence or absences (see Norberg et al. 2019), our novel contribution focuses on predicting species abundance. In practice, to obtain abundance estimates, both occupancy and abundance predictions should be combined (Denes et al. 2015).

We assessed whether a rescaling correction could improve the biases in abundance predictions between predicted and observed abundance. This bias appears systematically in quantitative ecological predictions (Pearce and Ferrier 2001, Fukaya et al. 2020, Ploton et al. 2020). We rescaled predicted values to take the range of observed values using the following formula: 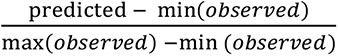 and assessed how this procedure affected model performance indicated by our evaluation metric set.

### Model cross-validations and transferability to novel climates

We evaluated model performance using two cross-validation strategies. We evaluated how well models predict abundance when i) interpolating within environments (within-sample) and ii) extrapolating into novel climate conditions (out-of-sample). The first scenario applies when models are interpolated to fill geographic gaps in sampling within a species range. The second scenario applies when modelling species abundance under climate change. When testing interpolation within-sample environments, we randomly held out 20% of the abundance data and fitted models to the remaining 80%. This within-sample model evaluation used a random subset of sites within the full covariate space.

Our second cross-validation strategy tested model transferability into novel conditions. Transferability measures if models can be projected beyond environments found within bounds of the covariate data. Given the rate of anthropogenic changes to our environment, models will be best applied when also accurate in novel conditions with no past analogues (Evans 2012, Sequeira et al. 2018a). Model transferability can be low if models are overfitted, exhibit non-stationarity, or are missing important covariates (Yates et al. 2018). We built separate models following the above protocol to test model transferability. To do so, we non-randomly sampled 20% data from above the 80^th^ quantile of sea-surface temperature in reef-fishes, and above the 80^th^ quantile of the climatological PCA-1 in birds, and fitted our abundance models to the remaining 80%. We estimated all evaluation metrics within the out-of-sample cross-validation sets as above. In both scenarios, we assumed that cross-validation frames were independent of the training data frames (Randin et al. 2006, Roberts et al. 2017).

We did not perform k-fold cross-validation for the full span of covariate space because we wanted to gain an understanding of abundance estimates from directional environmental novelty due to climate change (e.g., predicting abundance in warmer temperatures than fishes currently experience in the oceans). As a hypothetical example, if we split a temperature gradient from 20-30°C into 20-22, 22-24, 24-26, 26-28 and 28-30°C bins and examined performance on each bin; spatial auto-correlation would lead to an underestimate of model performance in novel future climates when evaluating the middle bins. Under temperature warming, we therefore only used the highest 20% bin threshold for exploring extrapolation (i.e., transferability to novel climates). To ensure cross-validation scenarios of interpolation and extrapolation were comparable, we used only one 20% subsample for the interpolation (random) subset also. Although this procedure is not encouraged in general for SDM fitting and evaluation, for good reason (Roberts et al. 2017), it suits our specific cross-validation goals (Sequeira et al. 2018a, Yates et al. 2018). We expected our findings to be robust to any small biases introduced by only performing one-fold cross-validations because of the high number of species included in the exercise. We did, however, perform 10-fold cross-validations when sub-sampling species absences to ensure findings were robust to variation in the locations of species absences.

### Species’ and data characteristics

We also tested how characteristics of species’ abundance, frequency, and data availability affected model performance. To explain variation in model performance among species, we calculated i) the mean abundance of species when present; ii) the proportion of presence compared to absence records (within 1000km) in the observational data (% occupied sites); and iii) the total number of presence records per species (overall observation number). Although the frequency of occurrence and the total number of presences are colinear in bird and fish datasets (rho=0.87, rho=0.67, respectively) we included both because unbiased estimates of coefficients are achieved through multiple regression (Morrissey and Ruxton 2018). We log10 transformed and standardised predictor variables to have unit variance and removed outliers (points > 2 SD from the mean) from the response variables. Next, we fitted multiple regressions that explained how the model evaluation metrics depended on our three measures of species’ characteristics. For simplicity, we present these results using D_spearman_ due to the high number of comparisons and the importance of model discrimination highlighted above. We first fitted a full model, including three two-way interactions between pairs of predictors. We performed backwards stepwise model selection and selected the model with the lowest AIC score using the R package MuMIn (Burnham and Anderson 2002, Barton 2017). We plotted marginal effects by predicting model effects for a given variable across the mean value of all other model covariates. We fitted these models using phylogenetic generalised least squares using the R package caper using maximum likelihood to estimate Pagels λ (Blomberg and Symonds 2014, Orme et al. 2018). We used published bird (Jetz et al. 2012, 2014; https://birdtree.org/downloads/) and fish (Rabosky et al. 2018; https://fishtreeoflife.org/downloads/) phylogenetic trees.

## Results

### Overview of model performance

We first assessed performance by applying all frameworks to all species and evaluating interpolative prediction of within-sample observations. Doing so, model performance was highly variable and generally low (Table S3; Figure S5-S8). For example, across all models and species, D_spearman_ had a median of 0.29 (5^th^ percentile = -0.17, 95^th^ percentile = 0.64), median D_slope_ was 0.06 (-0.07 – 0.47) and median A_mae_ was 0.74 (0.48 – 1.52). As such, of the complete model set (n=68), only 51% of species had at least one model with a D_spearman_ above 0.5; 53% of species had at least one model with a D_slope_ between 0.5 and 1.5, but 93% of species had at least one model with A_mae_ predicting mean abundances within 10% of observed mean abundances, and 33% of species had models fitting all the above criteria.

We next investigated the best fitting algorithm for each species independently, keeping only the single best model each species (i.e., our ‘optimal model’). Random forests were most often selected as the optimal models for discrimination (precision, accuracy) being best for 51% (55%, 46%) of the species, gradient boosting machines for 22% (26%, 32%) and generalised linear models and generalised additive models for 16% (9%, 14%) and 12% (10%, 8%) of species respectively (Figure 2, Figure S5, Figure S7, Figure S9). Building models using abundance-absence data led to the best discrimination (precision, accuracy) performance for 68% (30%, 26%) of species, 19% (24%, 32%) using only species’ abundance, and 14% (46%, 42%) using a two-step indirect approach relating abundance to occurrence probability (Figure S4).

**Figure 2.**
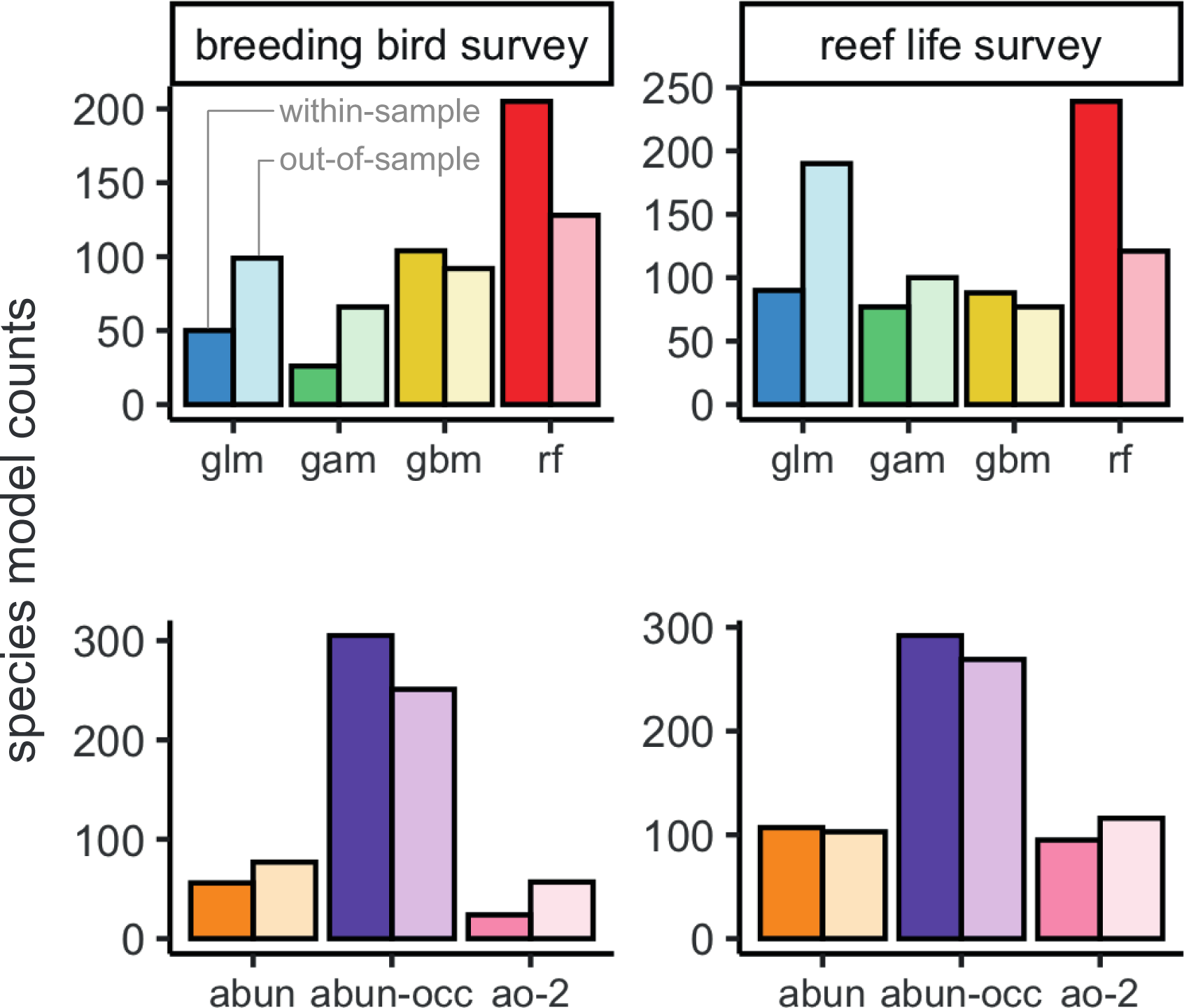
Counts of the model framework (top row) and abundance response treatment (bottom row) to which the most discriminatory model for each species belongs. Breeding bird survey shown in left column and reef life survey in right column. Colour shading indicates whether model predictions were from the within-sample model runs (dark) or out-of-sample model runs (light). See Figure S4 for counts using most accurate, most precise models, as well as combining all metric groups.

When selecting an optimal model for each species, model performance was good for most metrics (Figure 3; Table 2). For example, there were positive correlations for most species between observed and predicted abundances and the error of average abundance estimation was relatively low. Specifically, median D_spearman_ was 0.48 (0.14 – 0.72) and 0.43 (0.17 – 0.72) for bird and fish surveys respectively, and median A_mae_ was 0.62 (0.43 – 0.97) and 0.69 (0.46 – 1.34) respectively. Some measures of model performance were poor, leading to a biased relationship between observed and predicted abundances and a poor estimation of abundance variation. Specifically, D_slope_ was 0.25 (0.02 – 0.68) and 0.20 (0.01 – 0.99) for bird and fish surveys, and P_dispersion_ was 0.51 (0.12 – 1.27) and 0.44 (0.05 – 1.67), respectively.

**Figure 3.**
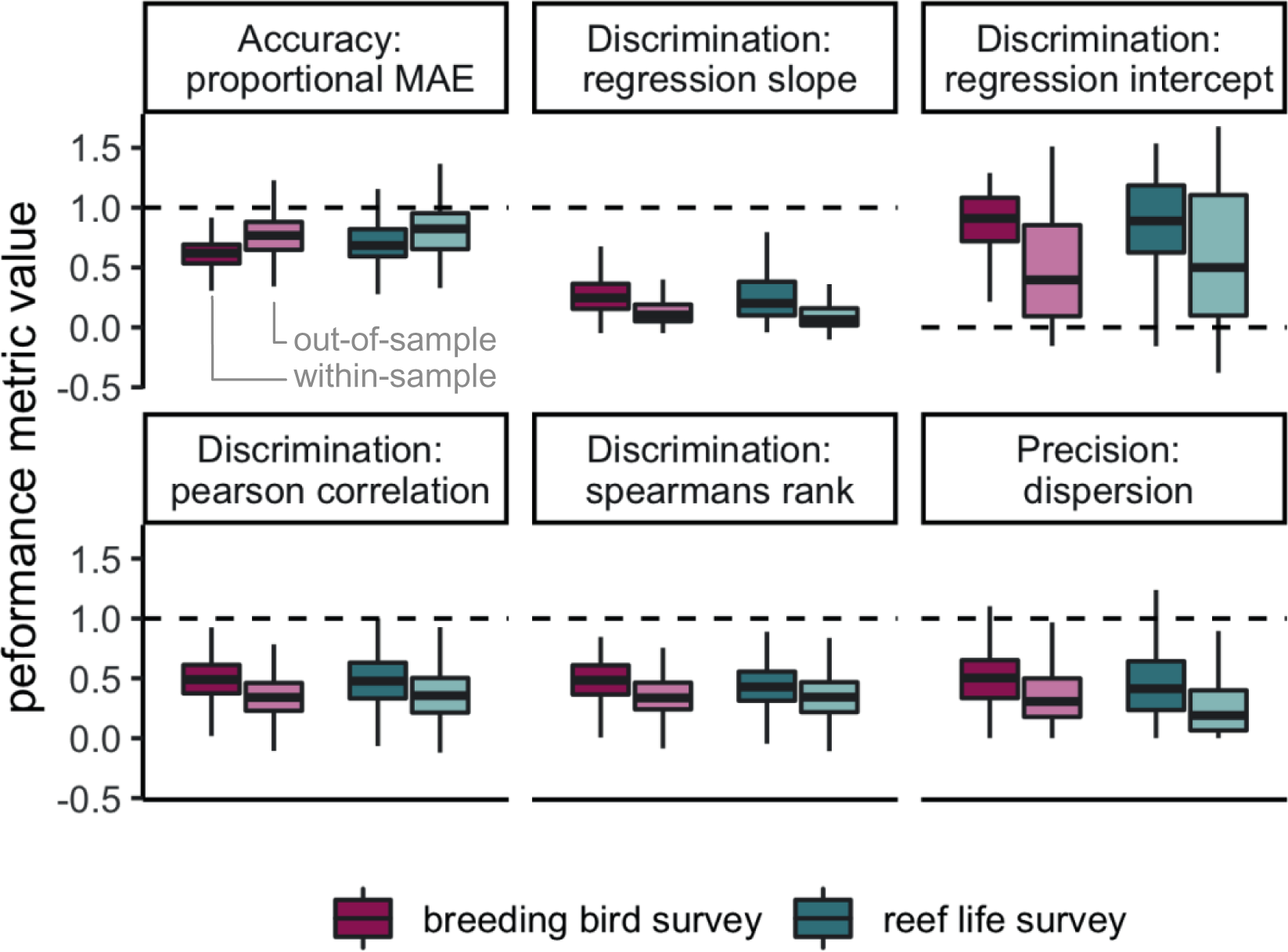
Boxplots of model performance of most discriminatory model for each species across all 6 metrics. Colours indicate breeding bird survey and reef life survey, whereas shading indicates within-sample and out-of-sample cross validations. Dashed lines indicate target values for each metric. Note that the type of model is not necessarily the same for a given species in the within-sample and out-of-sample comparisons, as indicated in Figure 2. Central lines correspond to median values, hinges correspond to 25^th^ and 75^th^ quantiles and whiskers correspond to 1.5x the hinges. Outliers are excluded from visualisations. See Figure S25 for performance of most accurate and most precise models, as well as combining all metric groups.

Predictions of abundance from optimal models had a high correspondence with observed abundances, on average across all species, in both fish and birds (Figure 4). However, as indicated by the evaluation metrics, the overall relationship was biased to be shallower than a 1:1 correspondence between observed and predicted abundance by models consistently overestimating low abundance and underestimating high abundances (Figure 4; see Figure S15 and S17 for all models, and Figure S19 and S21 for individual optimal models). Applying a rescaling correction (rescaling predicted abundances to the observed abundance range) for each species helped to correct this systematic bias. Model performance improved as indicated by A_mae_ (before correction = 0.64-0.69 to after correction = 0.88-0.94), D_slope_ (0.20-0.25 to 0.50-0.56) and P_dispersion_ (0.44-0.51 to 1.10), however, performance decreased when indicated by D_intercept_ (1.7-2.2 to 5.6-5.9; see full results in Table S4).

**Figure 4.**
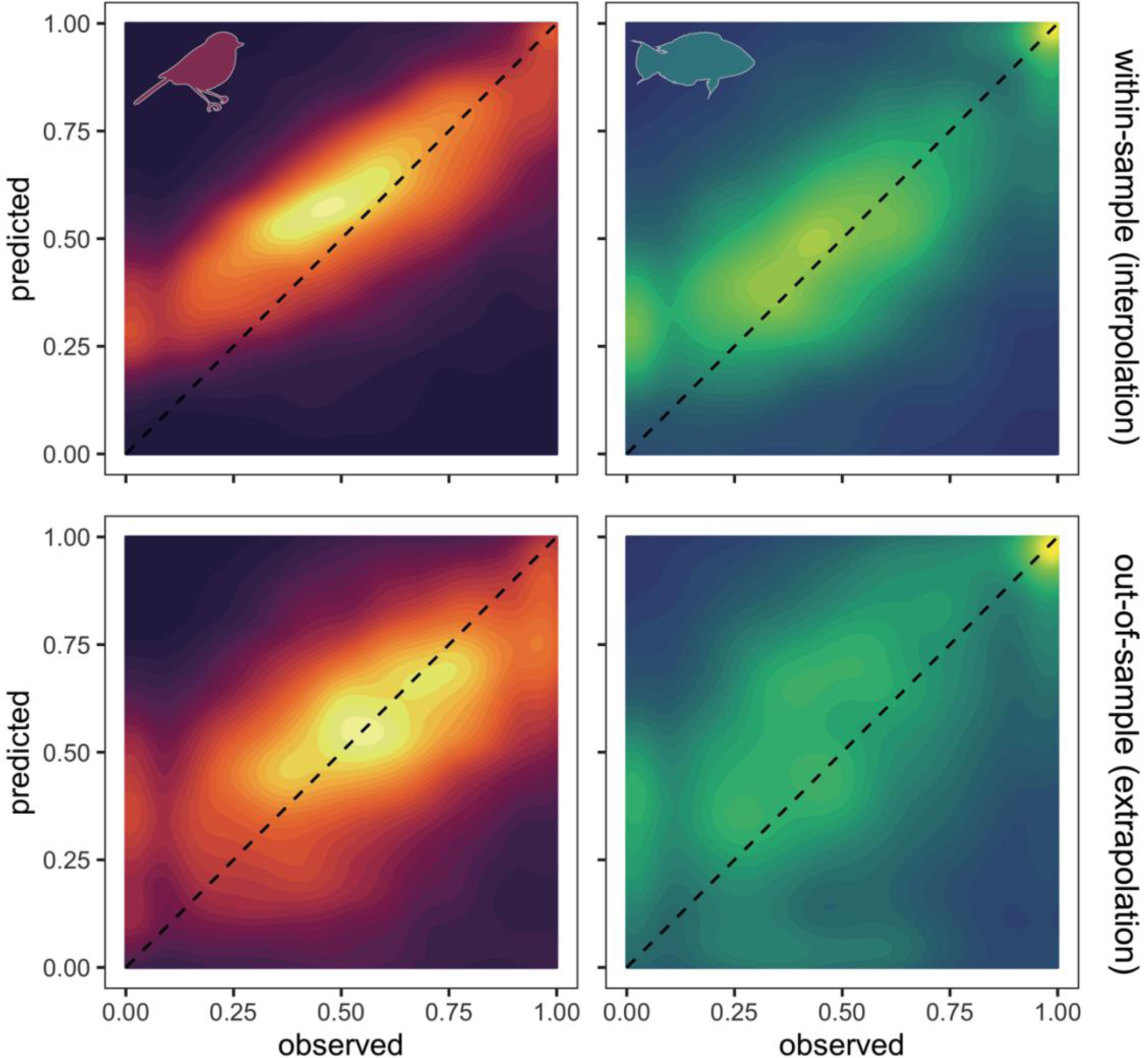
Contour plots of observed abundance vs. model predicted abundance across bird and fish datasets. Upper panels show within-sample interpolation and lower panels show out-of-sample extrapolation of predicted values (see Methods and Materials for details). Dashed line indicates 1:1 correspondence. Colour intensity indicates the number of records within contour. Both axes are log10+1 transformed and rescaled between 0 and 1 to show ability of models to discriminate abundance values. To avoid species with more data dominating patterns, for each species, we binned observations into 30 bins and estimated the mean predicted abundance for each observed abundance bin. Note that, due to the 0-1 transformation, a value of 0 is the minimum observed or predicted abundance value.

### Model transferability to novel conditions (i.e., out-of-sample)

Transferring models to novel conditions, the best performing algorithm for each species in terms of discrimination (precision, accuracy) shifted to generalised linear models being the best for 33% species (27%, 21%), random forests for 29% (40%, 38%), generalised additive models for 19% (19%, 21%), and gradient boosting machines for 19% (15%, 21%) of the species (Figure 2, Figure S4, S6, S8, S10). Building models using abundance-absence data remained the best performing treatment of response data in terms of discrimination (precision, accuracy) for 60% (33%, 25%) of species, with 21% (41%, 20%) of species having best models when using only species’ abundances, and 20% (26%, 56%) using a two-step approach (Figure 2).

Transferring models to novel conditions reduced model performance for most metrics across both birds and fishes (Table 2; Figure 3). The general discrimination of high and low abundances remained (median D_spearman_ was 0.34 for birds and 0.34 for fishes). D_slope_ declined by more than half compared to within-sample cross-validations (median D_slope_ was 0.10 for birds and 0.06 for fishes). Surprisingly, accuracy increased compared to within-sample cross-validations with a median of 0.78 and 0.83 in birds and fishes, respectively.

Predicted abundance still corresponded with observed abundances on average across all species, in both fishes and birds (Figure 4), despite the poorer model performance. However, similar issues with a biased intercept and slope exist in the out-of-sample cross-validations as for the within-sample cross-validations, and were similarly corrected for by the rescaling procedure (Figure 4; see Figure S16 and S18 for all models, and Figure S20 and S22 for individual optimal models; see Table S4 for comparisons with rescaling).

### Species’ and data characteristics

The variation in model performance explained by species and data characteristics varied among performance metrics, and was higher in general for within-sample (R^2^ = 0.04 – 0.44) compared to out-of-sample cross-validations (R^2^ = 0.01 – 0.33; Table S5 – S8). All six evaluation metrics were affected by species or data characteristics in both birds and fishes (Table S5 – S8). D_intercept_ had the most variation explained by species and data characteristics in both birds and fishes (R^2^ of 0.42-0.44).

We present the example metric D_spearman_, which had a R^2^ between 0.16 and 0.33. The effects of species and data characteristics on D_spearman_ were highly consistent across within and out-of-sample predictions and across both datasets (Figure 5; Figure S23; Table S5-8). More observations decreased the D_spearman_. Higher frequency of occurrence increased D_spearman_ but only if species also had high number of observations. Species with higher abundance had higher D_spearman_ only if species had high frequency too. This last effect was not evident for fish species in out-of-sample predictions. Phylogenetic signal (Pagel’s λ) in the residuals was very weak ranging from 0 to 0.17.

**Figure 5.**
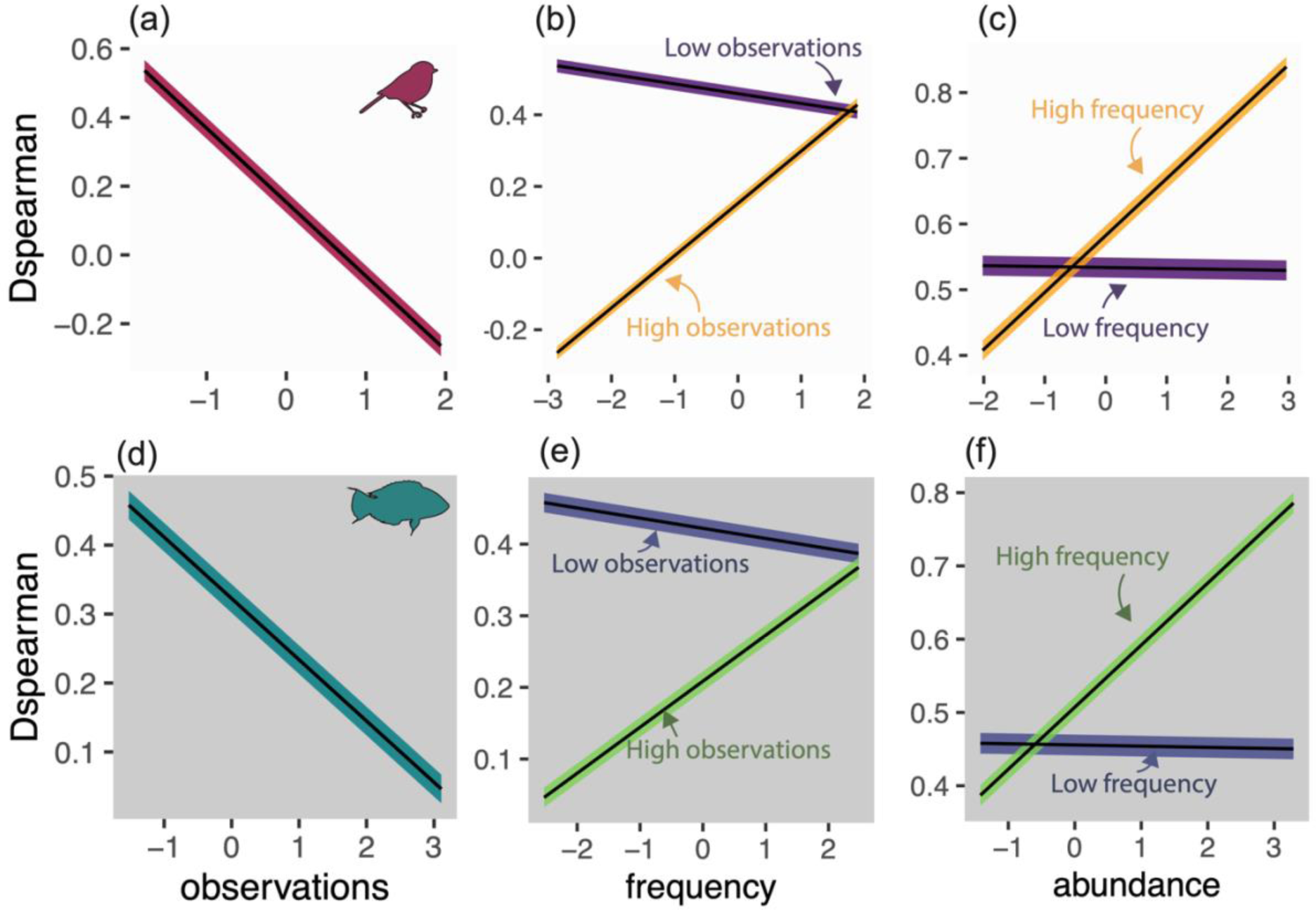
Effect of species’ and data characteristics on D_spearman_ for breeding bird survey (a-c) and reef life survey (d-f). Plots display marginal effects from multiple regressions fitted using phylogenetic generalised least squares for within-sample cross validations. Lines represent mean predicted values. Shaded areas show uncertainty as mean ± (standard error x 1.96) of coefficient values. All effects are significant at an alpha of 0.05, and interaction terms are only shown when significant. Full statistical results across all metrics, datasets and cross validations are displayed in (Table S5 to S8). See Figure S23 for effect of species and data characteristics on D_spearman_ in out-of-sample predictions.

## Discussion

We demonstrated the capacity to predict spatial patterns in abundance for many species if an appropriate model framework is chosen. The predictability of abundance using only the environmental response shapes of species has probably been under-appreciated somewhat, in part due to many options for statistical models and only a few providing acceptable predictions. For example, using GAMs and GLMs, Johnston et al. (2013) found a low rank correlation of 0.19 for predicted and observed seabird densities, and therefore focussed on coarser spatial scales for predictive analyses (also see (Illan et al. 2014)). Our results support that correlative abundance models could have an important role in quantifying the changing spatial patterns of species’ abundance due to environmental change, although many challenges remain. Here we discuss our relative success and failures in modelling abundance to better guide future applications.

### Successful aspects of species abundance models

A small number of good approaches for predicting species abundance emerged after exploring a large set of models. Correlation values from our optimal models were higher than ∼0.3 for more than 75% of species, and higher than ∼0.6 for 25% of species (Table 2). Our finding that random forests performed well at within-sample prediction provides solid evidence that the findings for Balearic shearwaters (*Puffinus mauretanicus*; Oppel et al. 2012) apply more generally, at least across the 800 species of bird and fish tested here. The high discrimination, precision and accuracy of random forests would improve confidence in assigning regions as important abundance-priority areas for conservation.

A focus on linear functions relating environments to local abundances may have previously reduced predictive performance. More flexible response curves of machine learning approaches allow for what may often be highly non-linear abundance niche shapes (Pearce and Ferrier 2001, Potts and Elith 2006, Renwick et al. 2012, Betts et al. 2019). Further optimised algorithms and deep learning approaches may better integrate abundance into biodiversity indicator frameworks given the much better performance of machine learning approaches here (Jetz et al. 2019). If abundance has been perceived to be poorly explained by climate or other variables in the past, it could be falsely concluded that broad-scale variables only weakly affect abundance and that abundance niches are more strongly constrained by factors other than species’ fundamental niches (but see Illan et al. 2014, Dallas and Santini 2020).

Accurate prediction of local abundances with abiotic variables supports the theoretical prediction that fitness optima along abiotic niche axes filters down to determine ecologically successful locations of high population growth rates (Maguire, 1973). The prediction of abundance from abiotic niche axes has been questioned by recent empirical studies (Dallas and Hastings 2018, Santini et al. 2019, Sporbert et al. 2020). These studies determine environmental effects on abundance indirectly from habitat suitability or environmental centroids. Here we directly relate abundance to environmental conditions which provides a more direct quantification of species’ abundance niche with fewer assumptions (Osorio-Olvera et al. 2020).

Our finding that modelling abundance directly was better than an indirect approach (i.e., comparing our abundance-absence models to two-stage models) for more than 80% of species indicates that spatial abundance and occurrence patterns are somewhat mismatched, or at least not always congruent (although it is challenging to completely disentangle abundance from occurrence, and vice versa). Mismatches arise from different ecological controls of abundance and occurrence, such as different demographic rates controlling each to different extents (McGill 2012, Johnston et al. 2015, Acevedo et al. 2017, Dallas and Santini 2020, Schulz et al. 2020, Yancovitch et al. 2020, Bohner and Diez 2020). Understanding such mismatches offers an important avenue for better understanding range and abundance shifts under climate change (Geppert et al. 2020) and potentially guiding spatial management and conservation. For example, a focus on occurrence can miss critical patches of high abundance driven by a few isolated factors (Johnston et al. 2015, Suggitt et al. 2018). Such ‘strongholds’ for species could be a common feature of ecological communities and are likely only considered when management is focussed on small scales for data-rich species. Moving species distribution models beyond modelling occurrences, to help identify such areas, will require improving knowledge of species’ responses to environmental gradients using multiple performance metrics (i.e., occurrence, abundance, demographic rates) (Ehrlén and Morris 2015, Ashcroft et al. 2017, Bohner and Diez 2020).

### Current limitations and challenges in species abundance models

We identify two important biases in abundance models here: why do we systematically over-predict low observed abundances and under-predict high observed abundances (see also Pearce and Ferrier 2001, Fukaya et al. 2020, Ploton et al. 2020)? And, why does having more abundance observations for a species lead to lower discriminatory power of predictions (i.e., poorer ability to discriminate between high abundance sites and low abundance sites)? These biases may jointly arise as we undoubtedly miss key biotic (e.g. ecological interactions) and micro-climatic variables from our models (Lembrechts et al. 2019), leading to extreme local abundances.

Missing inter- and intra-specific interactions has been a well-recognised problem in predictive occurrence-based species distribution modelling (Guisan and Thuiller 2005, Wisz et al. 2013, Mouquet et al. 2015, Pollock et al. 2020). For abundance-based models, species’ interactions can drive population feedbacks that may be important for explaining extreme abundances, but are missing from models in general, leading to poor predictive performance. Recent theoretical work highlights how interaction feedbacks can strongly modify abundance along environmental gradients, even if the fundamental niche shape is unimodal (Kéfi et al. 2016, Liautaud et al. 2019). In addition, behavioural aggregations from seasonal migrations or resource booms can lead to extreme abundances; challenging the identification of appropriate statistical response distributions (Lindén and Mäntyniemi 2011). These points emphasise the need to better understand how local environments, individual behaviour and species interactions together shape macroecological abundance patterns. Novel joint species distribution modelling approaches (Ovaskainen et al. 2017), or direct estimation of interaction strengths (Wootton and Emmerson 2005) are promising tools to help address such questions.

Abundance-based species distribution models could be further improved by considering fine-scale microclimatic data, a concept gaining traction for occurrence-based species distribution models (Potter et al. 2013, Bennie et al. 2014, Lembrechts et al. 2019) and critical for better conservation planning in the face of climate change (Roslin et al. 2009, Isaak et al. 2017). Microclimate variation within grid cells can arise from variations in topography, aspect (Bennie et al. 2008, Graae et al. 2018) and land-use features (Chen et al. 1999, 2006, Zhao et al. 2014, Senior et al. 2017) that filter species locally, and affect abundances, depending on species’ physiological and climatic niches (Ashcroft et al. 2014, Nowakowski et al. 2018, Waldock et al. 2020).

Incorporating (micro)climatic variation at the appropriate spatiotemporal scale for a given species is a critical area for model improvements (Roslin et al. 2009, Ashcroft et al. 2014, Rebaudo et al. 2016), especially for projections of future climate effects on species occurrence and abundance (Gillingham et al. 2012, Hannah et al. 2014, Maclean et al. 2015, Woods et al. 2015). Our sensitivity analysis indicates improved model fit with improved data resolution for some species, but not all, when using just one year of BBS data linked to a finer temporal resolution of climate data (Figure S25). This finding indicates species-specific behaviour (migratory vs. non-migratory), mobility (sedentary or mobile, home-range size), life-cycle (hibernators vs. year-round activity) and environmental niche characteristics (breadth, plasticity) could contribute to the resolution and windows of microclimatic data required to accurately estimate local abundances and occurrence (Bennie et al. 2014, Lembrechts et al. 2019).

An additional problem, not present in occurrence-based models, is that the probability of sampling a system in an extreme abundance state is higher with more samples, leading to outlier points (i.e., bright-spots or dark-spots). Perhaps these outliers could be an avenue to unveil important predictors of locations of hyper-abundance, or bright spots which in turn can comprise important targets for conservation (Cinner et al. 2016, Frei et al. 2018). Biased predictions and missed outliers have important consequences. For example, the shallower slope of predicted versus observed abundance will underestimate change in abundance when the environment changes. In contrast, the likelihood of persistence will be overestimated because abundance losses in the last stages of population decline are poorly captured by models such as ours (Bates et al. 2014). As such, separate models for occurrence and abundance patterns will need to be calibrated and outputs combined. For occurrence-based models more data generally leads to better models (Chefaoui et al. 2011), we identify the opposite here with the consequence that for abundance-based models data-poor species perhaps generate overconfident models, a caveat worth exploring further.

We identify that the transferability of species abundance models to novel environmental conditions is presently limited. This shortcoming applies to occurrence-based species distribution models (Sequeira et al. 2018a, Yates et al. 2018) and models of family-level abundances (Sequeira et al. 2018b), and may be exacerbated when considering species’ abundance. Models with perfect discrimination of presence-absence can still have poor predictive power of abundance values because more mechanisms underlie abundance variation and errors in capturing each mechanisms using statistical response functions will accumulate (Bahn and McGill 2013, Johnston et al. 2015). We demonstrate model performance also declines when predicting outside the bounds of even a single covariate (rather than a spatial block (Ploton et al. 2020)), with strong consequences for future climatic predictions.

Novel climatic conditions are fast emerging (Williams and Jackson 2007), hence solutions that improve model transferability are urgently needed (Radeloff et al. 2015, Harris et al. 2018). Whilst mechanistic models offer accurate predictions at coarse spatial scales (Fernandes et al. 2013, 2020), further integration with correlative frameworks may enable prediction at fine-scales and in novel environments (Cheung et al. 2008, Fernandes et al. 2020, Gamliel et al. 2020).

### Which species to target for abundance-based species distribution modelling?

Our consideration of strengths and limitations of species abundance models can help guide their application for predicting the spatial distribution of species abundance for systematic conservation planning (Margules and Pressey 2000, Pinsky et al. 2020, Pollock et al. 2020). Importantly, from a conservation perspective, we outline how model performance relates to rarity and thus extinction risk. Our results suggest that species with low frequency of occurrence and low mean abundance will be more challenging to predict. Perhaps such species are only weakly constrained by physiological niche limits, and more strongly constrained by meta-population dispersal, microclimate effects, and availability of resources, hosts, or prey items (Selig et al. 2014, Venter et al. 2014, Mouillot et al. 2016, Suggitt et al. 2018). In contrast, common and abundant species that mostly contribute to ecosystem functions and services may be good targets for species abundance modelling (Winfree et al. 2015, Mouillot et al. 2016). We also highlight how the treatment of abundance data can modify how well models perform in accuracy, discrimination and precision which could have important consequences depending on the target application (i.e., Figure S4). Here, consideration of species’ abundances as well as changes in occurrence should greatly assist understanding how biodiversity change affects ecosystem functioning and human wellbeing (Johnston et al. 2015, Kissling et al. 2018, Pinsky et al. 2020).

## Conclusions

Species’ abundances in localised field surveys can be predicted using broad-scale environmental and human factors, such as climate, land cover and habitat area for a large number of species. Species abundance models showed surprisingly similar performance in species from two very different ecological contexts. Transferring models to novel conditions was very challenging, however. Models fitted better for more frequently encountered and abundant species, highlighting that abundance models may be most applicable to questions relating to ecosystem function and service provision rather than in modelling rare or endemic species under extinction threats. When common species are to be prioritised (e.g., (Pinsky et al. 2020)), species abundance models could be used in many ways, providing spatial maps of species’ abundance, landscape scale estimates of ecological processes and services (Gilby et al. 2020), or helping to identify regions with large, stable, viable populations that can act as sources and facilitate reserve spill-over and ecosystem stability (Rondinini and Chiozza 2010, Halpern et al. 2010, Timus et al. 2017, Cabral et al. 2020; Table 1). We argue that spatial abundance models can provide critical biodiversity information with the potential to improve the ecological relevance and species conservation applications of species distribution models.

## Acknowledgements

We thank the many Reef Life Survey (RLS) divers who participated in data collection and provide ongoing expertise and commitment to the program. We thank the North American Breeding Bird Survey Dataset for providing access to the data and the thousands of participants who annually perform and coordinate the survey. CW was supported by the BiodivERsA grant Reef-Futures (no. 295340). JT acknowledges Research Council of Norway funded project BiodivERsA (Reef-Futures, no. 295340). Reef Life Survey uses the NCRIS-enabled Integrated Marine Observing System (IMOS) infrastructure for database support and storage, with support from Antonia Cooper and Elizabeth Oh. Thanks for IT and server support from Dominic Michel, Hussain Abbas and Benjamin Flück. Thanks to ‘Reef-Futures’ workshop attendees who provided constructive feedback on this work in addition to Jonathan Chase and three anonymous reviewers who provided valuable feedback on previous manuscript drafts.

## Notes

### Competing Interest Statement

The authors have declared no competing interest.

