## Appendix 1 for "A quantitative review of abundance-based species distribution models"

### Supporting tables

*Table S 1. Summary of gridded environmental variables used in all models.*

| Variable | Marine (M) or Terrestrial (T) | Spatial resolution | Transformation | Used in PCA | Source |
|---|---|---|---|---|---|
| Depth and elevation | M+T | 0.004° | none | no | GEBCO Compilation Group (2019)[1] |
| Chlorophyll-a (min, mean, max) | M | 0.25° | log10 | yes | Garnesson et al. (2019)[2] |
| Degree heating weeks (min, mean, max) | M | 0.25° | log10 | yes | Lui et al. (2014)[3] |
| Net primary productivity (min, mean, max) | M | 0.25° | log10 | yes | Behrenfeld et al. (1997)[4] |
| pH (min, mean, max) | M | 0.5° | none | yes | Norwegian Earth System Model forced ocean simulation (NorESM2) |
| Salinity (min, mean, max) | M | 0.25° | none | yes | Buongiorno Nardelli (2012)[5]; Droghei et al. (2016)[6]; Buongiorno Nardelli et al. (2016)[7] |
| Sea surface temperature (mean) | M | 0.25° | none | no | Lui et al. (2014)[3] |
| Coastal human population density (year 2015; sum in 50km$^2$) | M | 0.041° | log10 | no | Yeager, Marchand, Gill, Baum, & McPherson, (2017)[8] |
| Reef area (sum in 200km$^2$) | M | 0.041° | log10 | no | Yeager, Marchand, Gill, Baum, & McPherson, (2017)[8] |
| Wave energy (mean) | M | 0.041° | log10 | no | Yeager, Marchand, Gill, Baum, & McPherson, (2017)[8] |
| Annual Mean Temperature (bio1) | T | 0.083° | none | yes | MERRAclim[9] |
| Mean Diurnal Range Temperature (bio2) | T | 0.083° | none | yes | MERRAclim[9] |
| Isothermality (bio2/bio7) (* 100) (bio3) | T | 0.083° | none | yes | MERRAclim[9] |
| Temperature Seasonality (standard deviation *100) (bio4) | T | 0.083° | none | yes | MERRAclim[9] |
| Max Temperature of Warmest Month (bio5) | T | 0.083° | none | yes | MERRAclim[9] |
| Min Temperature of Coldest Month (bio6) | T | 0.083° | none | yes | MERRAclim[9] |
| Temperature Annual Range (bio5-bio6) (bio7) | T | 0.083° | none | yes | MERRAclim[9] |
| Mean temperature of most humid quarter (bio8) | T | 0.083° | none | yes | MERRAclim[9] |
| Mean temperature of least humid quarter (bio9) | T | 0.083° | none | yes | MERRAclim[9] |
| Mean temperature of warmest quarter (bio10) | T | 0.083° | none | yes | MERRAclim[9] |
| Mean temperature of coldest quarter (bio11) | T | 0.083° | none | yes | MERRAclim[9] |
| Annual mean specific humidity (bio12) | T | 0.083° | none | yes | MERRAclim[9] |
| Specific humidity of the most humid month (bio13) | T | 0.083° | none | yes | MERRAclim[9] |
| Specific humidity of the least humid month (bio14) | T | 0.083° | none | yes | MERRAclim[9] |
| Specific humidity seasonality (CV) (bio15) | T | 0.083° | none | yes | MERRAclim[9] |
| Specific humidity of most humid quarter (bio16) | T | 0.083° | none | yes | MERRAclim[9] |
| Specific humidity of least humid quarter (bio17) | T | 0.083° | none | yes | MERRAclim[9] |
| Specific humidity of warmest quarter (bio18) | T | 0.083° | none | yes | MERRAclim[9] |
| Specific humidity of colder quarter (bio19) | T | 0.083° | none | yes | MERRAclim[9] |
| Primary forest cover | T | 0.0083° | none | no | Hoskins et al. (2016)[10] |
| Human population density | T | 0.0416° | log10 | no | Gridded Population of the World v4[11] |

[1]GEBCO Compilation Group (2019) GEBCO 2019 Grid (doi:10.5285/836f016a-33be-6ddc-e053-6c86abc0788e) [2]Garnesson, P., Mangin, A., D'Andon, O. F., Demaria, J., & Bretagnon, M. (2019). The CMEMS GlobColour chlorophyll a product based on satellite observation: Multi-sensor merging and flagging strategies. Ocean Science, 15(3), 819–830. doi:10.5194/os-15-819-2019 [3]Liu, G., Heron, S. F., Mark Eakin, C., Muller-Karger, F. E., Vega-Rodriguez, M., Guild, L. S., … Lynds, S. (2014). Reef-scale thermal stress monitoring of coral ecosystems: New 5-km global products from NOAA coral reef watch. Remote Sensing, 6(11), 11579–11606. doi:10.3390/rs61111579 [4]Behrenfeld, M. J., & Falkowski, P. G. (1997). Photosynthetic rates derived from

satellite-based chlorophyll concentration. Limnology and Oceanography, 42(1), 1–20. doi:10.4319/lo.1997.42.1.0001 [5]Buongiorno Nardelli, B., 2012: A Novel Approach for the High-Resolution Interpolation of In Situ Sea Surface Salinity. J. Atmos. Ocean. Technol., 29, 867–879, doi:10.1175/JTECH-D-11-00099.1. [6]Droghei, R., B. Buongiorno Nardelli, and R. Santoleri, 2016: Combining in-situ and satellite observations to retrieve salinity and density at the ocean surface. J. Atmos. Oceanic Technol. doi:10.1175/JTECH-D-15-0194.1. [7]Buongiorno Nardelli, B., R. Droghei, and R. Santoleri, 2016: Multi-dimensional interpolation of SMOS sea surface salinity with surface temperature and in situ salinity data. Rem. Sens. Environ., doi:10.1016/j.rse.2015.12.052. [8]Yeager, L. A., Marchand, P., Gill, D. A., Baum, J. K., & McPherson, J. M. (2017). Marine Socio-Environmental Covariates: queryable: global layers of environmental and anthropogenic variables for marine ecosystem studies. Ecology, 98(7), 1976. doi:10.1002/ecy.1884 [9]Vega, G. C., Pertierra, L. R., & Olalla-Tárraga, M. Á. (2018). MERRAclim, a high-resolution global dataset of remotely sensed bioclimatic variables for ecological modelling. *Scientific Data*, *5*(1), 180070. doi:10.1038/sdata.2018.70 [10]Hoskins, A. J., Bush, A., Gilmore, J., Harwood, T., Hudson, L. N., Ware, C., … Ferrier, S. (2016). Downscaling land-use data to provide global 30'' estimates of five land-use classes. Ecology and Evolution, 6(9), 3040–3055. doi:10.1002/ece3.2104; [11] Center for International Earth Science Information Network - CIESIN - Columbia University. 2018. Gridded Population of the World, Version 4 (GPWv4): Population Density Adjusted to Match 2015 Revision UN WPP Country Totals, Revision 11. Palisades, NY: NASA Socioeconomic Data and Applications Center (SEDAC). https://doi.org/10.7927/H4F47M65. Accessed 16/04/2020 – population density adjusted to 2015.

Table S 2. Summary of models fitted and their characteristics.

| Response | Modelling framework | Transformation | Distribution | Zero-inflation |
|---|---|---|---|---|
| abundance-occupancy | linear model | log | gaussian | no |
| | | log10 | gaussian | |
| | generalised linear model | none | Poisson | no |
| | | | negative binomial | |
| | | | tweedie | |
| | | | Poisson | yes |
| | | | negative binomial | |
| | | | Tweedie | |
| | generalised additive model | log | gaussian | no |
| | | log10 | gaussian | |
| | | none | Poisson | |
| | | | negative binomial | |
| | | | tweedie | |
| | | | Poisson | yes |
| | random forests | none | regression tree | no |
| | | log | regression tree | |
| | | log-discrete | classification tree | |
| | | log10 | regression tree | |
| | | log10-discrete | classification tree | |
| | gradient boosting machines | log | gaussian | no |
| | | log-discrete | gaussian | |
| | | log10 | gaussian | |
| | | log10-discrete | gaussian | |
| | | none | Poisson | |
| abundance | linear model | log | gaussian | no |
| | | log10 | gaussian | |
| | generalised linear model | none | Poisson | no |
| | | | negative binomial | |
| | | | tweedie | |
| | generalised additive model | log | gaussian | no |
| | | log10 | gaussian | |
| | | none | Poisson | |
| | | | negative binomial | |
| | | | tweedie | |
| | random forests | none | regression tree | no |
| | | log | regression tree | |
| | | log-discrete | classification tree | |
| | | log10 | regression tree | |
| | | log10-discrete | classification tree | |
| | gradient boosting machines | log | gaussian | no |
| | | log-discrete | gaussian | |
| | | log10 | gaussian | |
| | | log10-discrete | gaussian | |
| | | none | Poisson | |
| abundance-2 stage | linear model | log | gaussian | no |
| | | log10 | gaussian | |
| | generalised linear model | none | Poisson | no |
| | | | negative binomial | |
| | | | tweedie | |
| | | | Poisson | yes |
| | | | negative binomial | |
| | | | tweedie | |
| | generalised additive model | log | gaussian | no |
| | | log10 | gaussian | |
| | | none | Poisson | |
| | | | negative binomial | |
| | | | tweedie | |
| | | | Poisson | yes |
| | random forests | none | regression tree | no |
| | | log | regression tree | |
| | | log-discrete | classification tree | |
| | | log10 | regression tree | |
| | | log10-discrete | classification tree | |
| | gradient boosting machines | log | gaussian | no |
| | | log-discrete | gaussian | |
| | | log10 | gaussian | |
| | | log10-discrete | gaussian | |
| | | none | Poisson | |

Table S 3. Evaluation metric summaries across all fitted models for all species (n = 206,869).

| | within-sample | | | | | out-of-sample | | | | |
|---|---|---|---|---|---|---|---|---|---|---|
| **metric** | **Q0.05** | **IQR0.25** | **median** | **IQR0.75** | **Q0.95** | **Q0.05** | **IQR0.25** | **median** | **IQR0.75** | **Q0.95** |
| $A_{mae}$ | 0.48 | 0.62 | 0.74 | 0.88 | 1.52 | 0.49 | 0.71 | 0.89 | 1.00 | 2.57 |
| $D_{intercept}$ | 0.74 | 1.69 | 3.29 | 7.43 | 38.34 | 0.01 | 0.72 | 2.63 | 7.65 | 50.68 |
| $D_{slope}$ | -0.07 | 0.01 | 0.06 | 0.17 | 0.47 | -0.16 | -0.01 | 0.00 | 0.05 | 0.26 |
| $D_{pearson}$ | -0.20 | 0.07 | 0.24 | 0.40 | 0.63 | -0.32 | -0.10 | 0.06 | 0.24 | 0.51 |
| $D_{spearman}$ | -0.17 | 0.12 | 0.29 | 0.44 | 0.64 | -0.35 | -0.08 | 0.10 | 0.27 | 0.51 |
| $P_{dispersion}$ | 0.04 | 0.18 | 0.34 | 0.58 | 1.51 | 0.00 | 0.05 | 0.18 | 0.44 | 1.88 |

*Table S 4. Summary metrics comparing raw and rescaled values for median, 25$^{th}$ and 75$^{th}$ percentiles of model performance values across all species.*

| | | | 25$^{th}$ percentile | | median | | 75th percentile | |
|---|---|---|---|---|---|---|---|---|
| **cross validation** | **dataset** | **metric** | **raw** | **rescale** | **raw** | **rescale** | **raw** | **rescale** |
| within-sample | breeding bird survey | $A_{mae}$ | 0.54 | 0.71 | 0.62 | 0.94 | 0.7 | 1.5 |
| | | $D_{intercept}$ | 1.4 | 3.1 | 2.2 | 5.9 | 3.4 | 10 |
| | | $D_{pearson}$ | 0.37 | 0.37 | 0.49 | 0.49 | 0.61 | 0.61 |
| | | $D_{slope}$ | 0.15 | 0.4 | 0.25 | 0.56 | 0.36 | 0.7 |
| | | $D_{spearman}$ | 0.36 | 0.36 | 0.48 | 0.48 | 0.61 | 0.61 |
| | | $P_{dispersion}$ | 0.34 | 0.94 | 0.51 | 1.1 | 0.68 | 1.4 |
| | reef life survey | $A_{mae}$ | 0.59 | 0.63 | 0.69 | 0.88 | 0.84 | 1.5 |
| | | $D_{intercept}$ | 0.9 | 2.3 | 1.7 | 5.6 | 5.7 | 20 |
| | | $D_{pearson}$ | 0.33 | 0.33 | 0.48 | 0.48 | 0.63 | 0.63 |
| | | $D_{slope}$ | 0.1 | 0.35 | 0.2 | 0.5 | 0.38 | 0.67 |
| | | $D_{spearman}$ | 0.31 | 0.31 | 0.43 | 0.43 | 0.56 | 0.56 |
| | | $P_{dispersion}$ | 0.25 | 0.9 | 0.44 | 1.1 | 0.72 | 1.3 |
| out-of-sample | breeding bird survey | $A_{mae}$ | 0.65 | 0.72 | 0.78 | 1.1 | 0.92 | 1.7 |
| | | $D_{intercept}$ | 0.4 | 2.4 | 1.6 | 4.8 | 4.4 | 11 |
| | | $D_{pearson}$ | 0.23 | 0.23 | 0.34 | 0.34 | 0.46 | 0.46 |
| | | $D_{slope}$ | 0.051 | 0.26 | 0.1 | 0.38 | 0.19 | 0.54 |
| | | $D_{spearman}$ | 0.24 | 0.24 | 0.34 | 0.34 | 0.46 | 0.46 |
| | | $P_{dispersion}$ | 0.18 | 0.96 | 0.32 | 1.1 | 0.55 | 1.4 |
| | reef life survey | $A_{mae}$ | 0.66 | 0.7 | 0.83 | 1.1 | 0.97 | 1.9 |
| | | $D_{intercept}$ | 0.42 | 2.4 | 1.5 | 6.9 | 4.7 | 27 |
| | | $D_{pearson}$ | 0.21 | 0.21 | 0.36 | 0.36 | 0.5 | 0.5 |
| | | $D_{slope}$ | 0.017 | 0.24 | 0.065 | 0.41 | 0.16 | 0.6 |
| | | $D_{spearman}$ | 0.22 | 0.22 | 0.34 | 0.34 | 0.47 | 0.47 |
| | | $P_{dispersion}$ | 0.071 | 0.96 | 0.2 | 1.2 | 0.41 | 1.4 |

*Table S 5. Model summary table for breeding bird survey within-sample interpolations, presenting coefficients and statistical tests of evaluation metrics values in response to species' mean abundance, frequency of occurrence and number of observations, accompanying Figure 5.*

| metric | data/species characteristic | coef | se | z-value | p-value | r-squared |
|---|---|---|---|---|---|---|
| Amae | abundance | 0.06 | 0.01 | 7.29 | <0.001 | 0.28 |
| Amae | frequency | -0.12 | 0.01 | -7.95 | <0.001 | 0.28 |
| Amae | observations | 0.08 | 0.01 | 5.58 | <0.001 | 0.28 |
| Amae | abundance:frequency | -0.02 | 0.01 | -3.09 | <0.01 | 0.28 |
| Amae | frequency:observations | -0.03 | 0.01 | -3.9 | <0.001 | 0.28 |
| Dintercept | abundance | 1.21 | 0.11 | 10.91 | <0.001 | 0.42 |
| Dintercept | frequency | 0.64 | 0.19 | 3.26 | <0.01 | 0.42 |
| Dintercept | observations | -0.36 | 0.18 | -1.95 | 0.05 | 0.42 |
| Dintercept | abundance:frequency | 0.65 | 0.17 | 3.79 | <0.001 | 0.42 |
| Dintercept | abundance:observations | -0.35 | 0.19 | -1.85 | 0.06 | 0.42 |
| Dslope | abundance | 0.04 | 0.01 | 3.89 | <0.001 | 0.19 |
| Dslope | frequency | 0.09 | 0.02 | 4.96 | <0.001 | 0.19 |
| Dslope | observations | -0.09 | 0.02 | -5.12 | <0.001 | 0.19 |
| Dslope | abundance:frequency | 0.05 | 0.02 | 3 | <0.01 | 0.19 |
| Dslope | abundance:observations | -0.03 | 0.02 | -1.89 | 0.06 | 0.19 |
| Dslope | frequency:observations | 0.02 | 0.01 | 1.9 | 0.06 | 0.19 |
| Dpearson | abundance | 0.05 | 0.01 | 5.76 | <0.001 | 0.31 |
| Dpearson | frequency | 0.11 | 0.02 | 6.28 | <0.001 | 0.31 |
| Dpearson | observations | -0.12 | 0.02 | -7.54 | <0.001 | 0.31 |
| Dpearson | abundance:frequency | 0.04 | 0.02 | 2.73 | <0.01 | 0.31 |
| Dpearson | abundance:observations | -0.03 | 0.02 | -1.88 | 0.06 | 0.31 |
| Dpearson | frequency:observations | 0.03 | 0.01 | 2.87 | <0.01 | 0.31 |
| Dspearman | abundance | 0.05 | 0.01 | 5.81 | <0.001 | 0.33 |
| Dspearman | frequency | 0.09 | 0.02 | 5.55 | <0.001 | 0.33 |
| Dspearman | observations | -0.08 | 0.02 | -5.23 | <0.001 | 0.33 |
| Dspearman | abundance:frequency | 0.02 | 0.01 | 2.38 | <0.05 | 0.33 |
| Dspearman | frequency:observations | 0.05 | 0.01 | 4.92 | <0.001 | 0.33 |
| Pdispersion | abundance | 0.02 | 0.02 | 1.08 | 0.28 | 0.07 |
| Pdispersion | frequency | 0.08 | 0.03 | 2.56 | <0.05 | 0.07 |
| Pdispersion | observations | -0.09 | 0.03 | -2.87 | <0.01 | 0.07 |
| Pdispersion | abundance:frequency | 0.1 | 0.03 | 3.59 | <0.001 | 0.07 |
| Pdispersion | abundance:observations | -0.07 | 0.03 | -2.24 | <0.05 | 0.07 |

*Table S 6. Model summary table for breeding bird survey out-of-sample extrapolations, presenting coefficients and statistical tests of evaluation metrics values in response to species' mean abundance, frequency of occurrence and number of observations, accompanying Figure S24.*

| metric | data/species characteristic | coef | se | z-value | p-value | r-squared |
|---|---|---|---|---|---|---|
| Amae | abundance | 0.07 | 0.02 | 4.6 | <0.001 | 0.11 |
| Amae | frequency | -0.04 | 0.03 | -1.26 | 0.21 | 0.11 |
| Amae | observations | 0.04 | 0.03 | 1.56 | 0.12 | 0.11 |
| Amae | abundance:frequency | -0.04 | 0.01 | -3.02 | <0.01 | 0.11 |
| Dintercept | abundance | 0.98 | 0.22 | 4.43 | <0.001 | 0.08 |
| Dintercept | frequency | -1.49 | 0.4 | -3.74 | <0.001 | 0.08 |
| Dintercept | observations | 1.16 | 0.38 | 3.06 | <0.01 | 0.08 |
| Dintercept | frequency:observations | -0.38 | 0.2 | -1.89 | 0.06 | 0.08 |
| Dslope | abundance | 0.02 | 0.01 | 2.34 | <0.05 | 0.05 |
| Dslope | observations | -0.01 | 0.01 | -2.12 | <0.05 | 0.05 |
| Dslope | abundance:observations | 0.02 | 0.01 | 2.51 | <0.05 | 0.05 |
| Dpearson | abundance | 0.05 | 0.01 | 5.14 | <0.001 | 0.17 |
| Dpearson | frequency | 0.03 | 0.02 | 1.72 | 0.09 | 0.17 |
| Dpearson | observations | -0.06 | 0.02 | -3.52 | <0.001 | 0.17 |
| Dpearson | abundance:frequency | 0.02 | 0.01 | 1.74 | 0.08 | 0.17 |
| Dpearson | frequency:observations | 0.02 | 0.01 | 2.19 | <0.05 | 0.17 |
| Dspearman | abundance | 0.05 | 0.01 | 5.66 | <0.001 | 0.23 |
| Dspearman | frequency | 0.06 | 0.02 | 3.59 | <0.001 | 0.23 |
| Dspearman | observations | -0.07 | 0.02 | -4.29 | <0.001 | 0.23 |
| Dspearman | abundance:frequency | 0.02 | 0.01 | 2.53 | <0.05 | 0.23 |
| Dspearman | frequency:observations | 0.03 | 0.01 | 2.84 | <0.01 | 0.23 |
| Pdispersion | observations | -0.03 | 0.02 | -1.87 | 0.06 | 0.01 |

*Table S 7. Model summary table for reef-life survey within-sample interpolations, presenting coefficients and statistical tests of evaluation metrics values in response to species' mean abundance, frequency of occurrence and number of observations, accompanying Figure 5.*

| **metric** | **data/species characteristic** | **coef** | **se** | **z-value** | **p-value** | **r-squared** |
|---|---|---|---|---|---|---|
| Amae | abundance | 0.1 | 0.01 | 10.56 | <0.001 | 0.24 |
| Amae | frequency | -0.02 | 0.01 | -1.8 | 0.07 | 0.24 |
| Amae | observations | 0.02 | 0.01 | 1.33 | 0.18 | 0.24 |
| Amae | abundance:frequency | -0.03 | 0.01 | -2.65 | <0.01 | 0.24 |
| Amae | abundance:observations | 0.03 | 0.01 | 2.28 | <0.05 | 0.24 |
| Dintercept | abundance | 15.01 | 0.87 | 17.22 | <0.001 | 0.44 |
| Dintercept | observations | -0.07 | 0.8 | -0.09 | 0.93 | 0.44 |
| Dintercept | abundance:observations | 2.91 | 0.82 | 3.54 | <0.001 | 0.44 |
| Dslope | abundance | 0.04 | 0.01 | 3.13 | <0.01 | 0.08 |
| Dslope | frequency | -0.02 | 0.02 | -1.2 | 0.23 | 0.08 |
| Dslope | observations | -0.05 | 0.02 | -2.31 | <0.05 | 0.08 |
| Dslope | abundance:frequency | 0.03 | 0.01 | 1.97 | <0.05 | 0.08 |
| Dslope | frequency:observations | 0.02 | 0.01 | 1.57 | 0.12 | 0.08 |
| Dpearson | abundance | 0.05 | 0.01 | 4.77 | <0.001 | 0.11 |
| Dpearson | frequency | 0 | 0.01 | 0.27 | 0.79 | 0.11 |
| Dpearson | observations | -0.05 | 0.01 | -3.68 | <0.001 | 0.11 |
| Dpearson | frequency:observations | 0.02 | 0.01 | 2.18 | <0.05 | 0.11 |
| Dspearman | abundance | 0.04 | 0.01 | 5.72 | <0.001 | 0.16 |
| Dspearman | frequency | 0.04 | 0.01 | 3.43 | <0.01 | 0.16 |
| Dspearman | observations | -0.05 | 0.01 | -4.35 | <0.001 | 0.16 |
| Dspearman | abundance:frequency | 0.02 | 0.01 | 2.39 | <0.05 | 0.16 |
| Dspearman | frequency:observations | 0.02 | 0.01 | 2.42 | <0.05 | 0.16 |
| Pdispersion | abundance | 0.04 | 0.02 | 1.49 | 0.14 | 0.04 |
| Pdispersion | frequency | -0.09 | 0.02 | -3.63 | <0.001 | 0.04 |
| Pdispersion | abundance:frequency | 0.05 | 0.02 | 2.22 | <0.05 | 0.04 |

*Table S 8. Model summary table for reef-life survey out-of-sample extrapolations, presenting coefficients and statistical tests of evaluation metrics values in response to species' mean abundance, frequency of occurrence and number of observations, accompanying Figure S24.*

| metric | data/species characteristic | coef | se | z-value | p-value | r-squared |
|---|---|---|---|---|---|---|
| Amae | abundance | 0.13 | 0.01 | 11.71 | <0.001 | 0.29 |
| Amae | frequency | -0.04 | 0.02 | -2.63 | <0.01 | 0.29 |
| Amae | observations | 0.05 | 0.02 | 3.01 | <0.01 | 0.29 |
| Amae | abundance:frequency | -0.03 | 0.02 | -2.05 | <0.05 | 0.29 |
| Amae | abundance:observations | 0.02 | 0.02 | 1.45 | 0.15 | 0.29 |
| Amae | frequency:observations | -0.02 | 0.01 | -1.94 | 0.05 | 0.29 |
| Dintercept | abundance | 14.31 | 1.13 | 12.72 | <0.001 | 0.33 |
| Dintercept | frequency | 1.36 | 1.63 | 0.84 | 0.4 | 0.33 |
| Dintercept | observations | -0.46 | 1.59 | -0.29 | 0.77 | 0.33 |
| Dintercept | abundance:frequency | 6.67 | 1.58 | 4.23 | <0.001 | 0.33 |
| Dintercept | abundance:observations | -4.05 | 1.5 | -2.7 | <0.01 | 0.33 |
| Dslope | abundance | -0.01 | 0.01 | -1.58 | 0.11 | 0.05 |
| Dslope | observations | -0.02 | 0.01 | -3.93 | <0.001 | 0.05 |
| Dpearson | abundance | 0.02 | 0.01 | 1.88 | 0.06 | 0.11 |
| Dpearson | frequency | 0.03 | 0.01 | 1.85 | 0.07 | 0.11 |
| Dpearson | observations | -0.08 | 0.01 | -5.85 | <0.001 | 0.11 |
| Dspearman | frequency | 0.02 | 0.01 | 1.28 | 0.2 | 0.05 |
| Dspearman | observations | -0.05 | 0.01 | -3.77 | <0.001 | 0.05 |
| Dspearman | frequency:observations | 0.02 | 0.01 | 2 | <0.05 | 0.05 |
| Pdispersion | abundance | -0.04 | 0.01 | -2.46 | <0.05 | 0.04 |
| Pdispersion | observations | -0.04 | 0.01 | -2.56 | <0.05 | 0.04 |

### Supporting figures

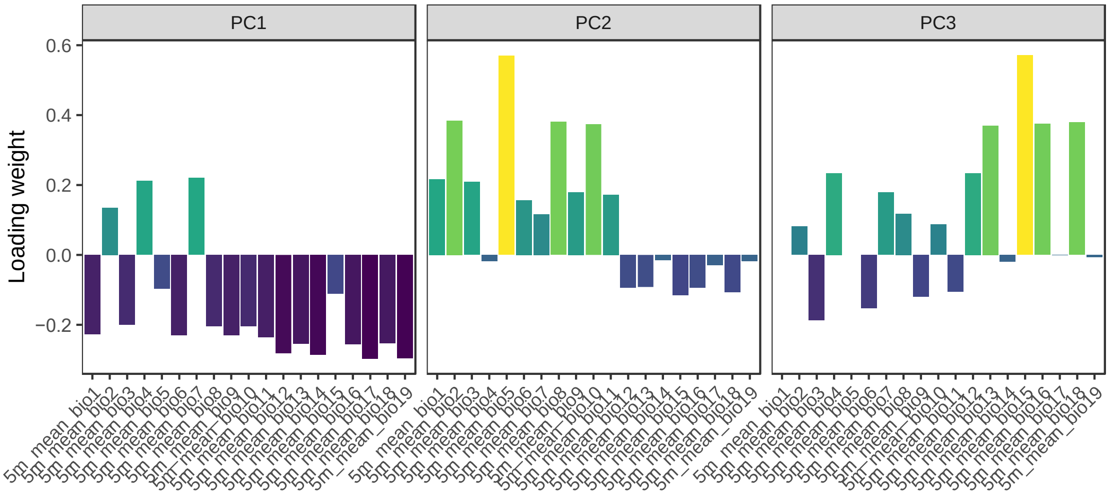


*Figure S1. Robust-PCA factor loadings representing the correlations between variables and principal components 1-3 for breeding bird survey climatological variables.*

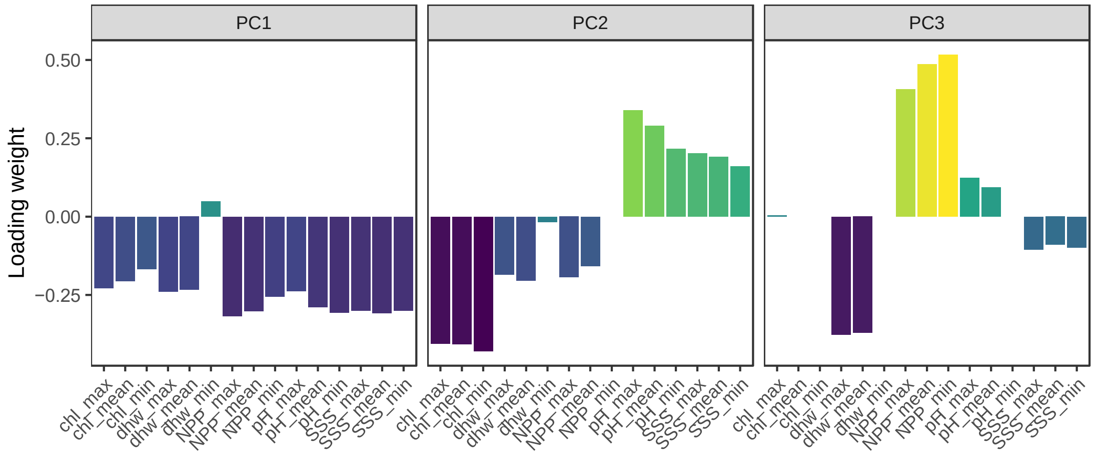


*Figure S2. Robust-PCA factor loadings representing the correlations between variables and principal components 1-3 for reef life survey climatological variables.*

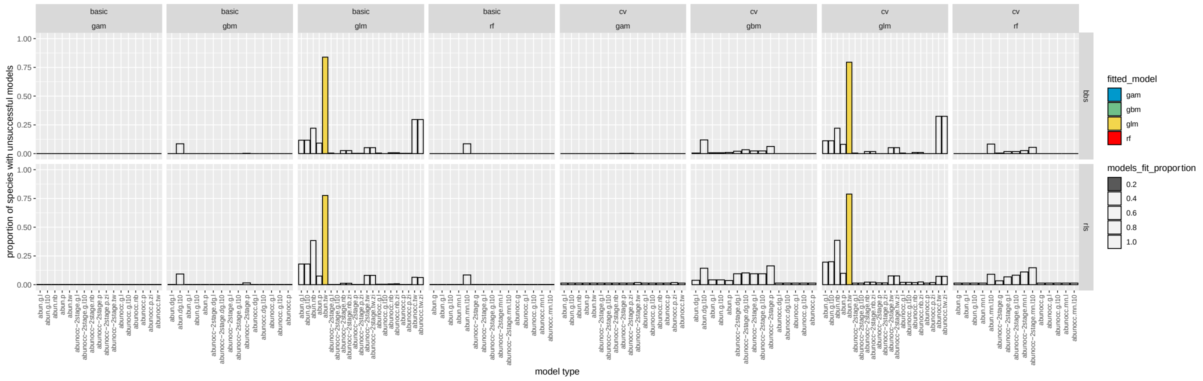


*Figure S3. Proportion of models unsuccessfully fitted across cross validations, datasets and modelling frameworks. Colour indicates modelling framework, and darker shading indicates a higher proportion of model fits were unsuccessful. Most models were fitted for all species but multinomial-log10 transformed models were sometimes unsuccessful, likely where low abundance makes transformations impossible.*

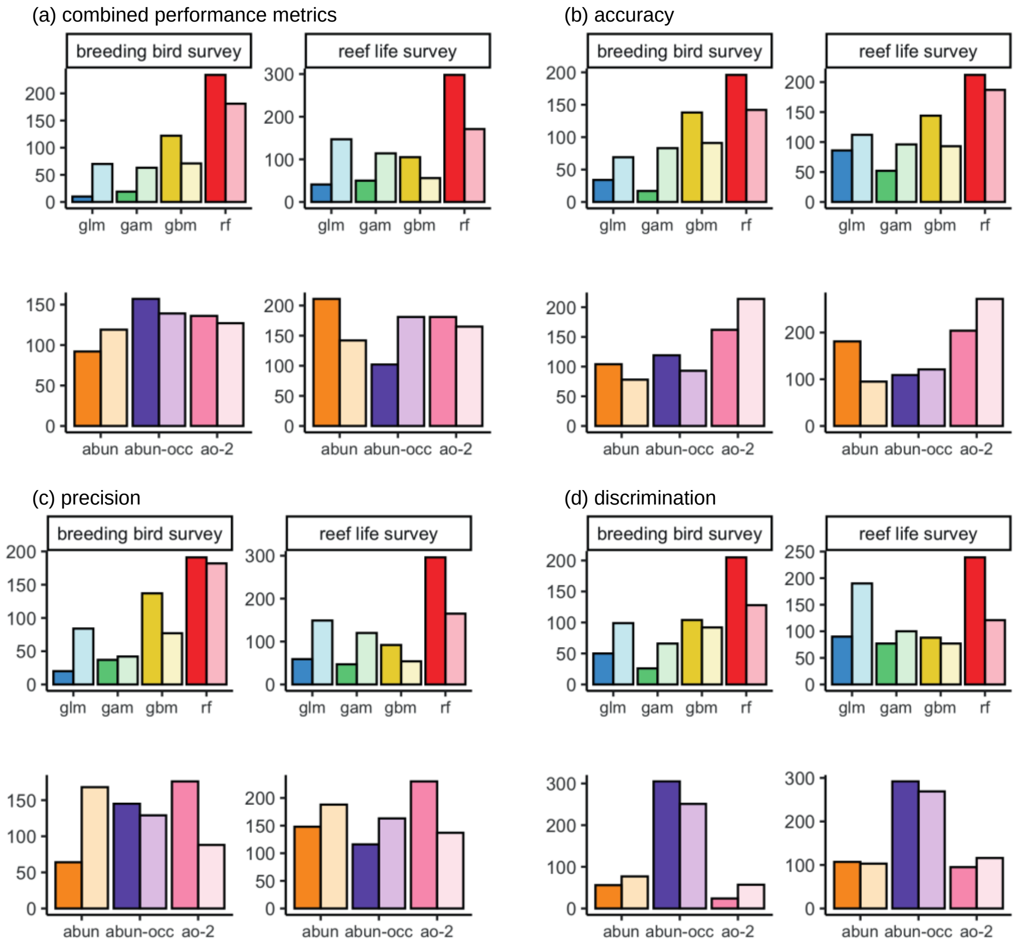


*Figure S4.* Counts of the most discriminatory model for each species showing in which model framework (top row) and abundance response treatment (bottom row) each optimal model belongs, across breeding bird survey (left column) and reef life survey (right column) for: (a) all performance metrics combined, (b) accuracy, (c) precision, (d) discrimination. Colour shading indicates whether model predictions were from the within-sample model runs (dark) or out-of-sample model runs (light).

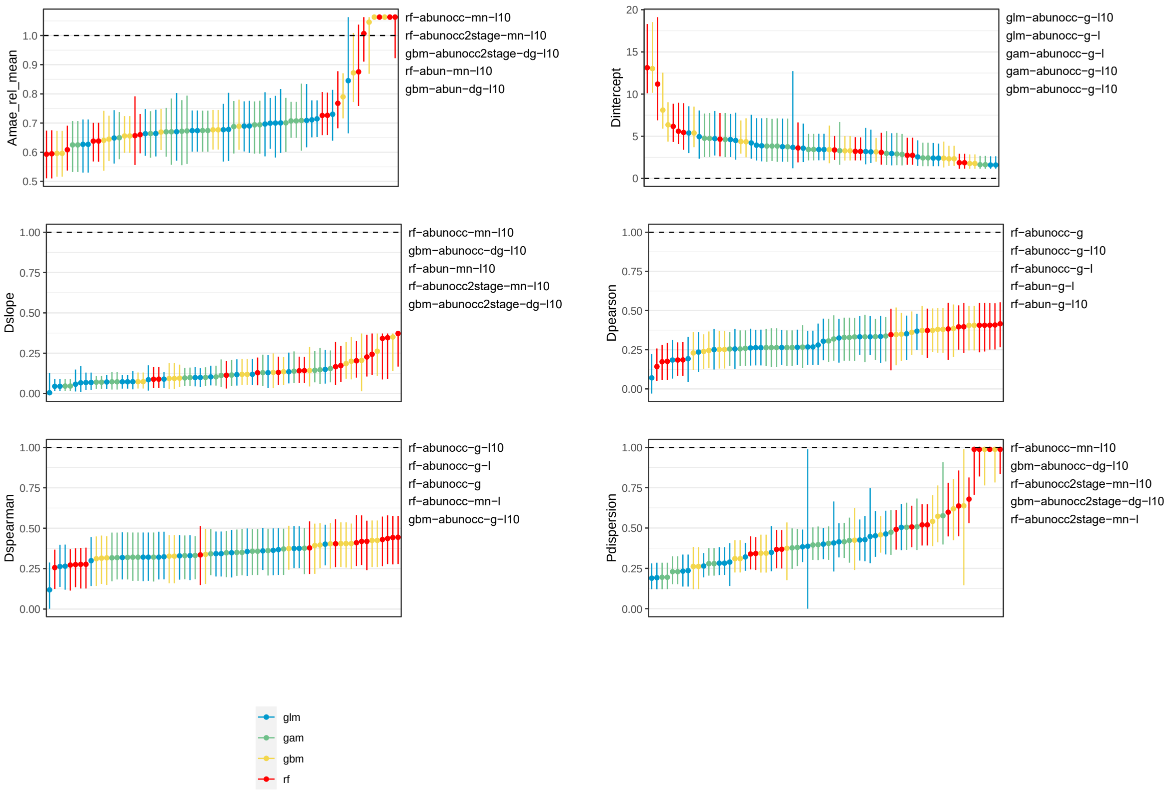


*Figure S5. Model evaluation metrics for breeding bird survey interpolated (i.e., within-sample) cross validations. X-axis is organised with increasing model performance from left-to-right. Dashed line indicates target for metrics. Best performing models are shown as text labels on the right of panels. Colour indicates modelling framework. We excluded extreme values exceeding the $10^{th}$ quantile away from target values for visualisation. Points indicate median species values within a modelling framework and lines are the $25^{th}$ to $75^{th}$ quantiles.*

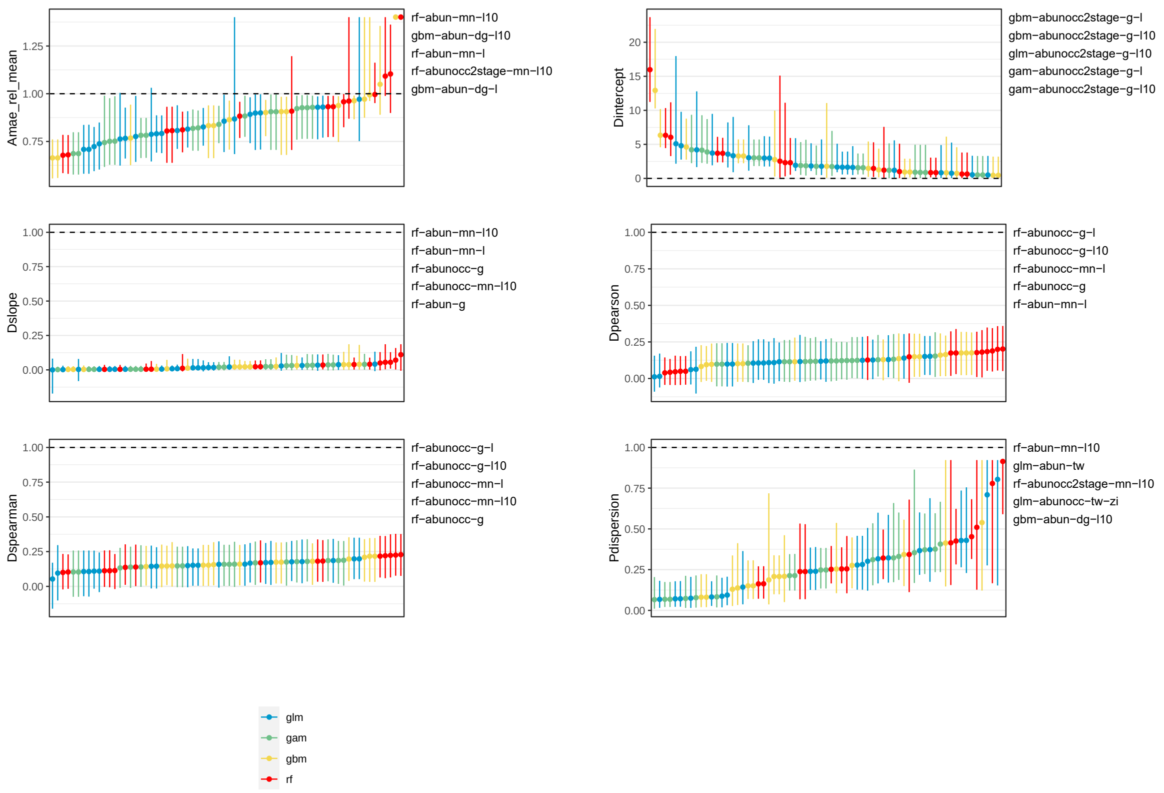


*Figure S6. Model evaluation metrics for breeding bird survey extrapolated (i.e., out-of-sample) cross validations. X-axis is organised with increasing model performance from left-to-right. Dashed line indicates target for metrics. Best performing models are shown as text labels on the right of panels. Colour indicates modelling framework. We excluded extreme values exceeding the 10$^{th}$ quantile away from target values for visualisation. Points indicate median species values within a modelling framework and lines are the 25$^{th}$ to 75$^{th}$ quantiles.*

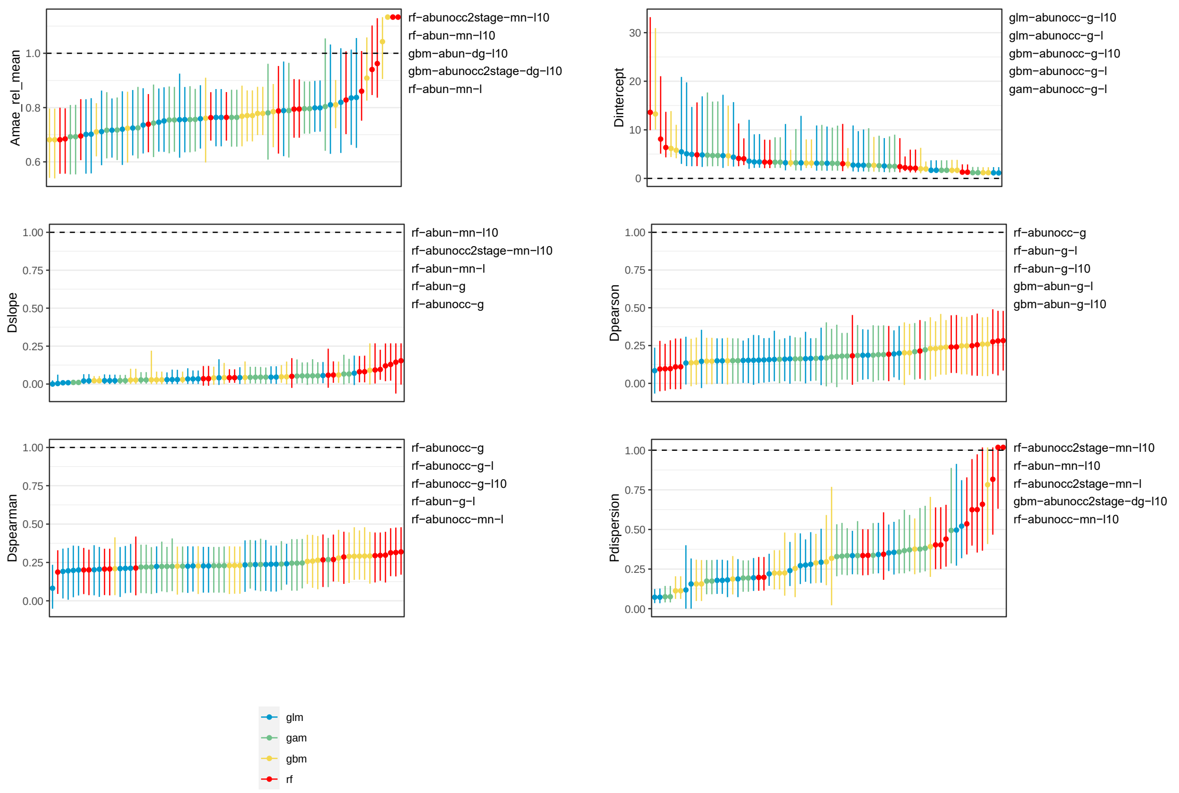


*Figure S7. Model evaluation metrics for reef life survey interpolated (i.e., within-sample) cross validations. X-axis is organised with increasing model performance from left-to-right. Dashed line indicates target for metrics. Best performing models are shown as text labels on the right of panels. Colour indicates modelling framework. We excluded extreme values exceeding the 10$^{th}$ quantile away from target values for visualisation. Points indicate median species values within a modelling framework and lines are the 25$^{th}$ to 75$^{th}$ quantiles.*

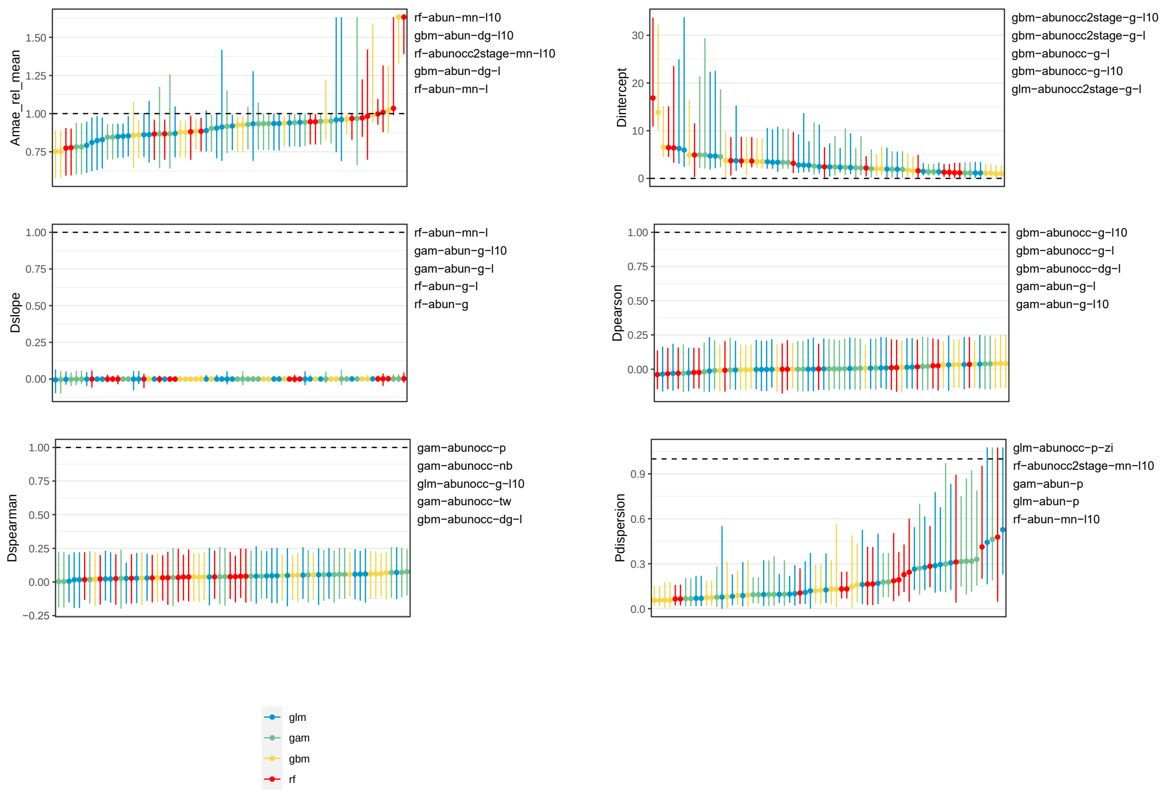


*Figure S8. Model evaluation metrics for reef life survey extrapolated (i.e., out-of-sample) cross validations. X-axis is organised with increasing model performance from left-to-right. Best performing models are shown as text labels on the right of panels. Colour indicates modelling framework. Dashed line indicates target for metrics. We excluded extreme values exceeding the 10$^{th}$ quantile away from target values for visualisation. Points indicate median species values within a modelling framework and line ranges are the 25$^{th}$ to 75$^{th}$ quantiles.*

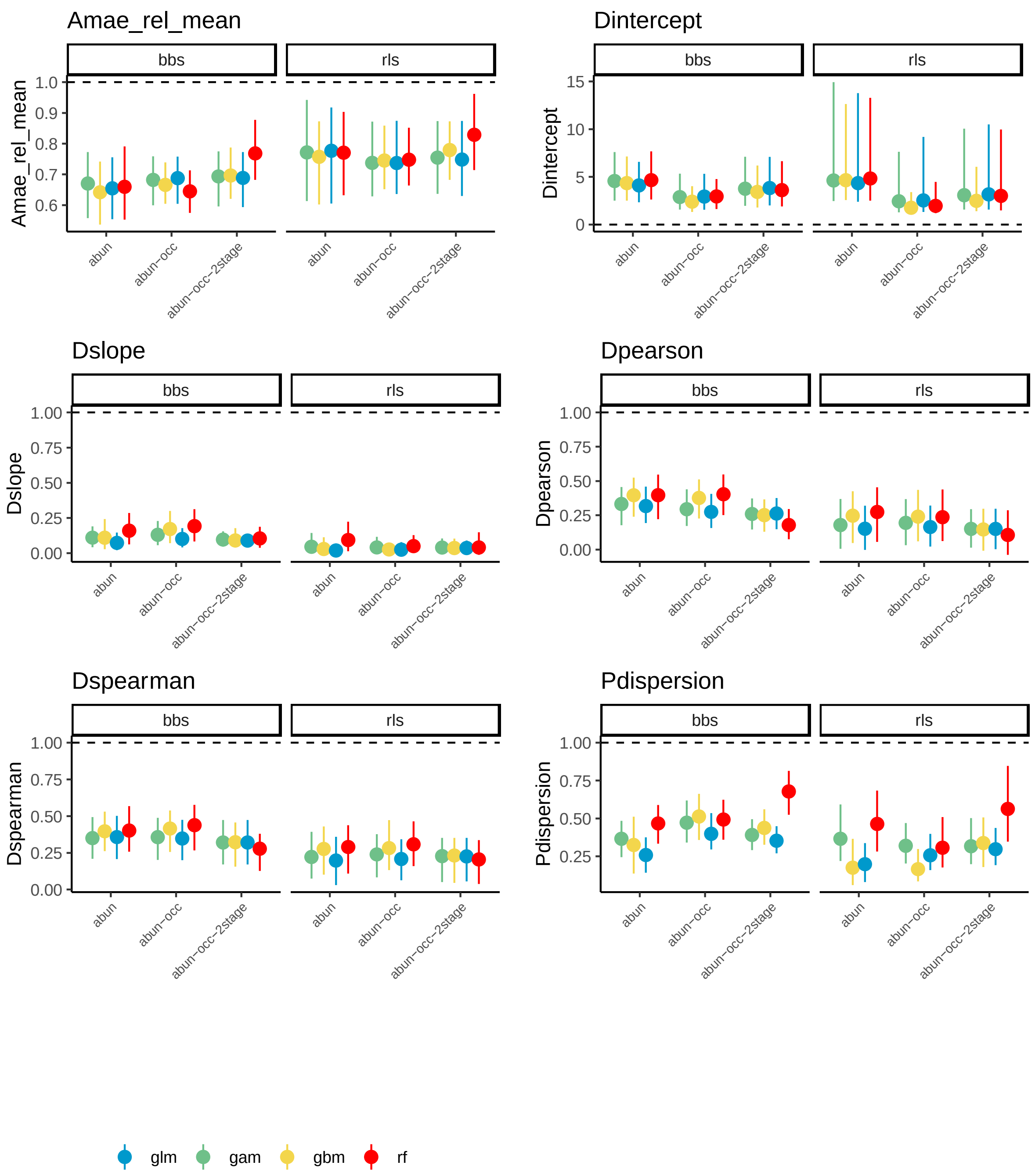


*Figure S9. Model evaluation metrics of across both reef life survey (rls) and breeding bird survey (bbs) for interpolated predictions (i.e., within-sample) aggregated to modelling frameworks and abundance types. Abundance types are abundance-only (abun), abundance and occurrence (abunocc) and 2stage species abundance models (abunocc-2stage), see methods for details. Points are median values across models and species and line ranges represent 25$^{th}$ to 75$^{th}$ quantiles. Colour indicates modelling framework. Dashed line indicates target for metrics.*

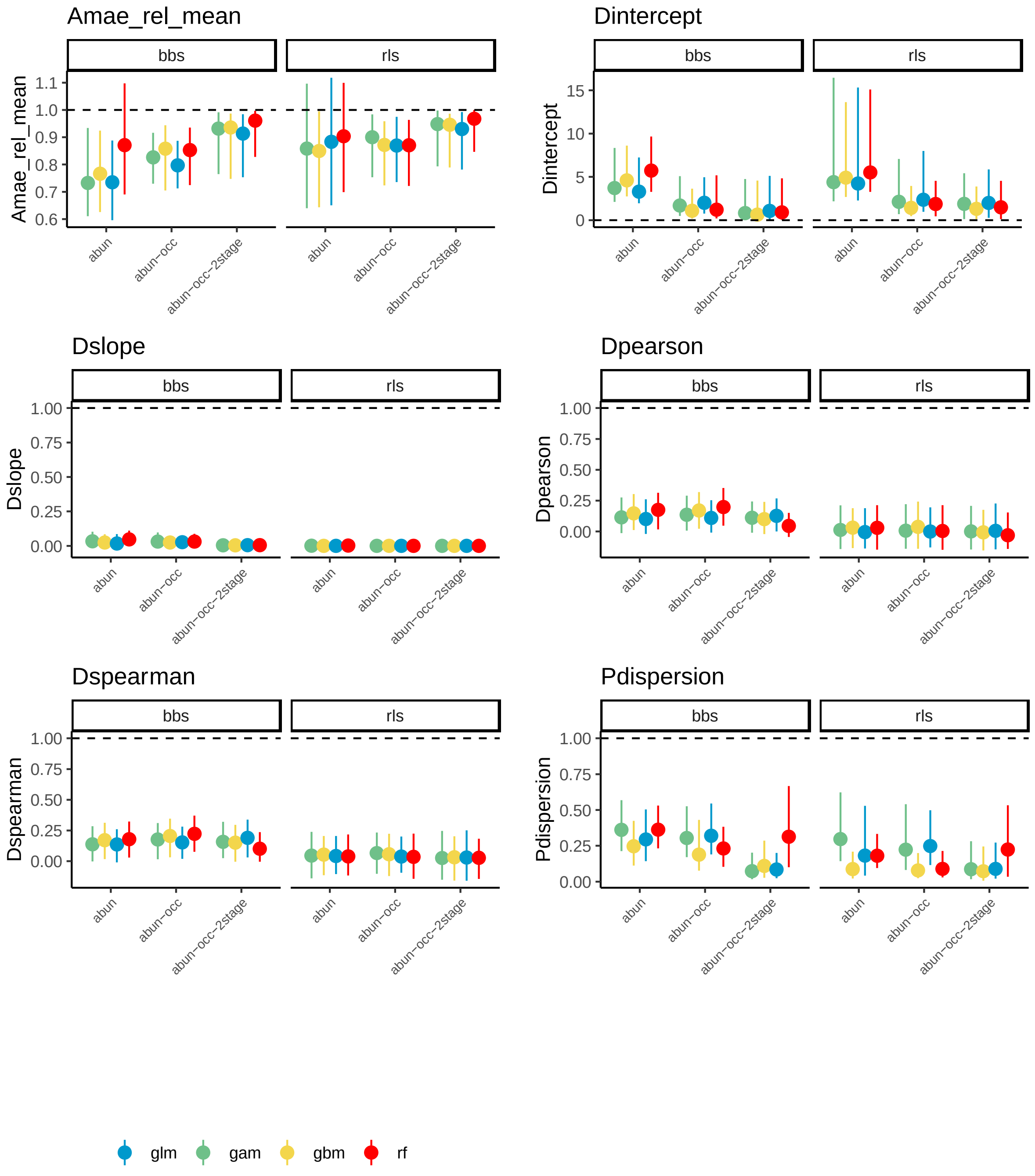


*Figure S10. Model evaluation metrics of across both reef life survey (rls) and breeding bird survey (bbs) for extrapolated predictions (i.e., out-of-sample) aggregated to modelling frameworks and abundance types. Abundance types are abundance-only (abun), abundance and occurrence (abunocc) and 2stage species abundance models (abunocc-2stage), see methods for details. Points are median values across models and species and line ranges represent 25$^{th}$ to 75$^{th}$ quantiles. Colour indicates modelling framework. Dashed line indicates target for metrics.*

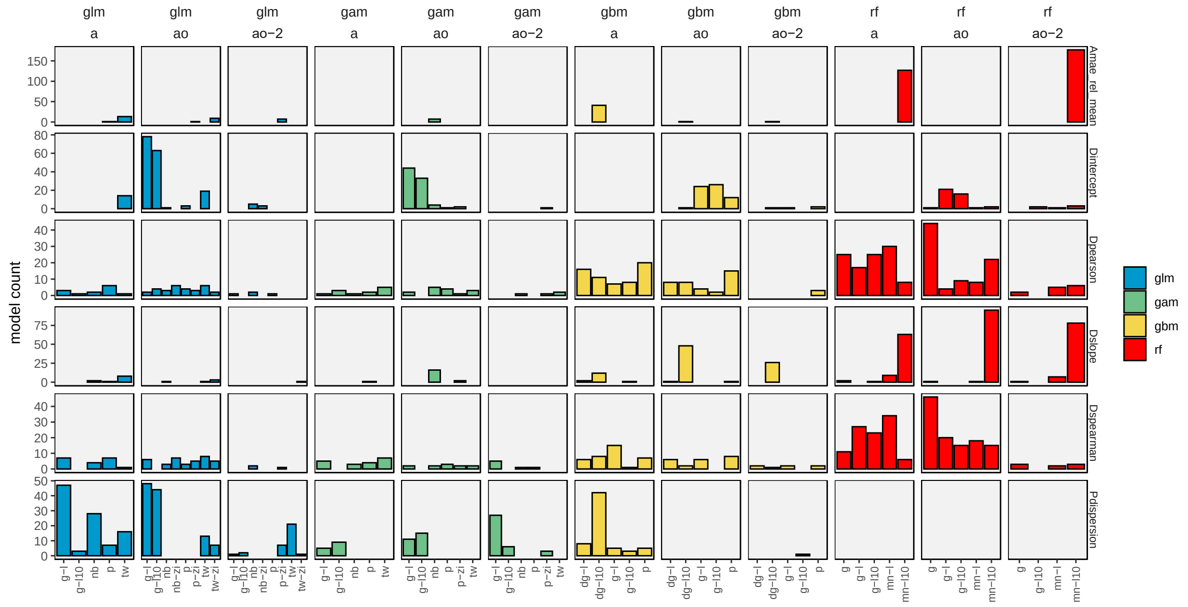


*Figure S11. Count of best models across all breeding bird survey species in interpolated projections (i.e., within-sample). X-axis shows different transformation and statistical error distributions categories, and column panels show different modelling frameworks and response data. Row panels show different evaluation metrics. Colour indicates modelling framework.*

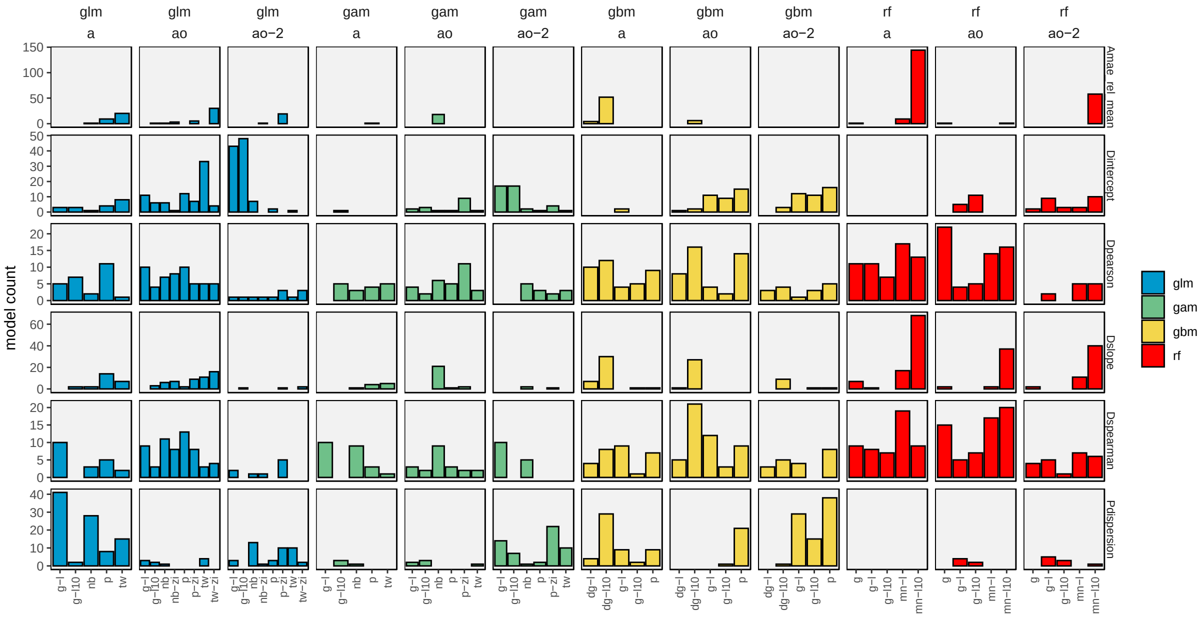


*Figure S12. Count of best models across all breeding bird survey species in extrapolated projections (i.e., out-of-sample). X-axis shows different transformation and statistical error distributions categories, and column panels show different modelling frameworks and response data. Row panels show different evaluation metrics. Colour indicates modelling framework.*

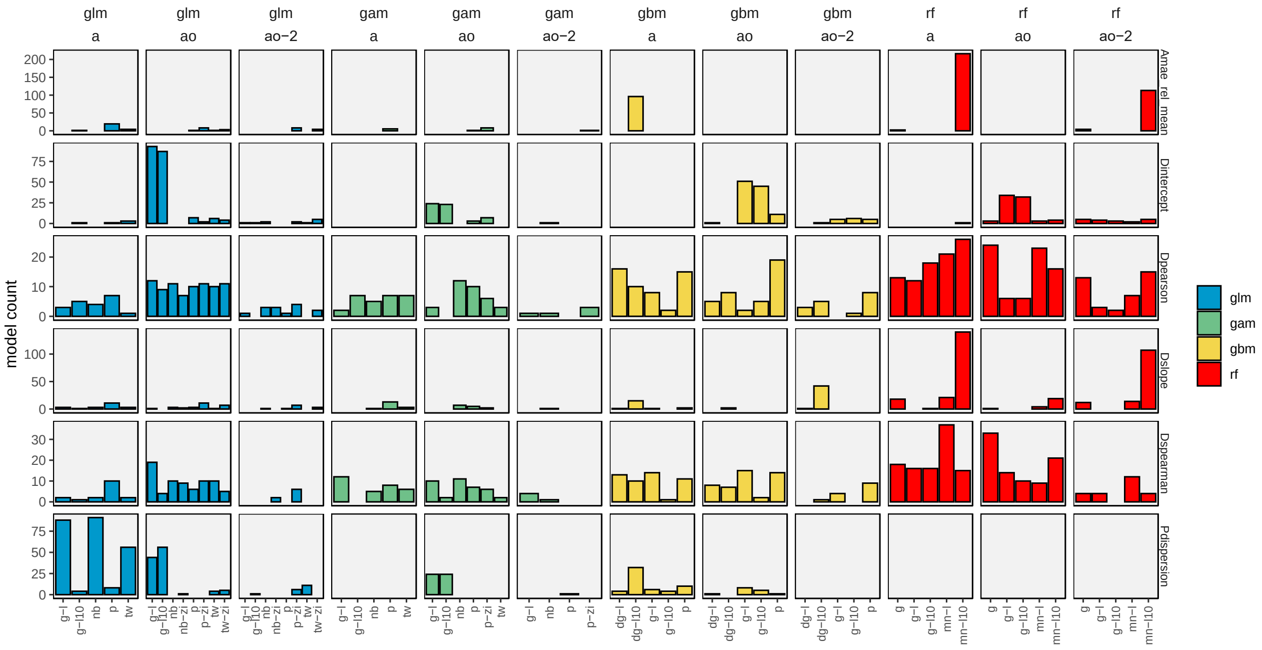


*Figure S13. Count of best models across all reef life survey species in interpolated projections (i.e., within-sample). X-axis shows different transformation and statistical error distributions categories, and column panels show different modelling frameworks and response data. Row panels show different evaluation metrics. Colour indicates modelling framework.*

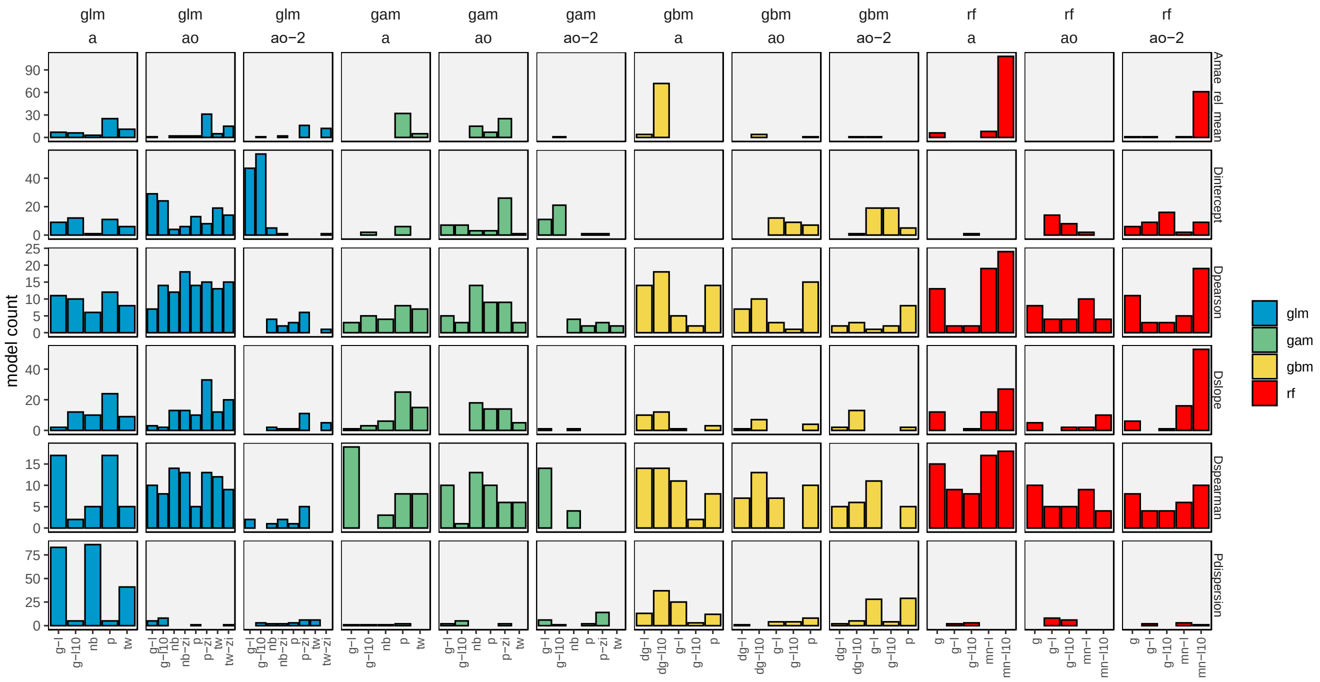


*Figure S14. Count of best models across all reef life survey species in extrapolated projections (i.e., out-of-sample). X-axis shows different transformation and statistical error distributions categories, and column panels show different modelling frameworks and response data. Row panels show different evaluation metrics. Colour indicates modelling framework.*

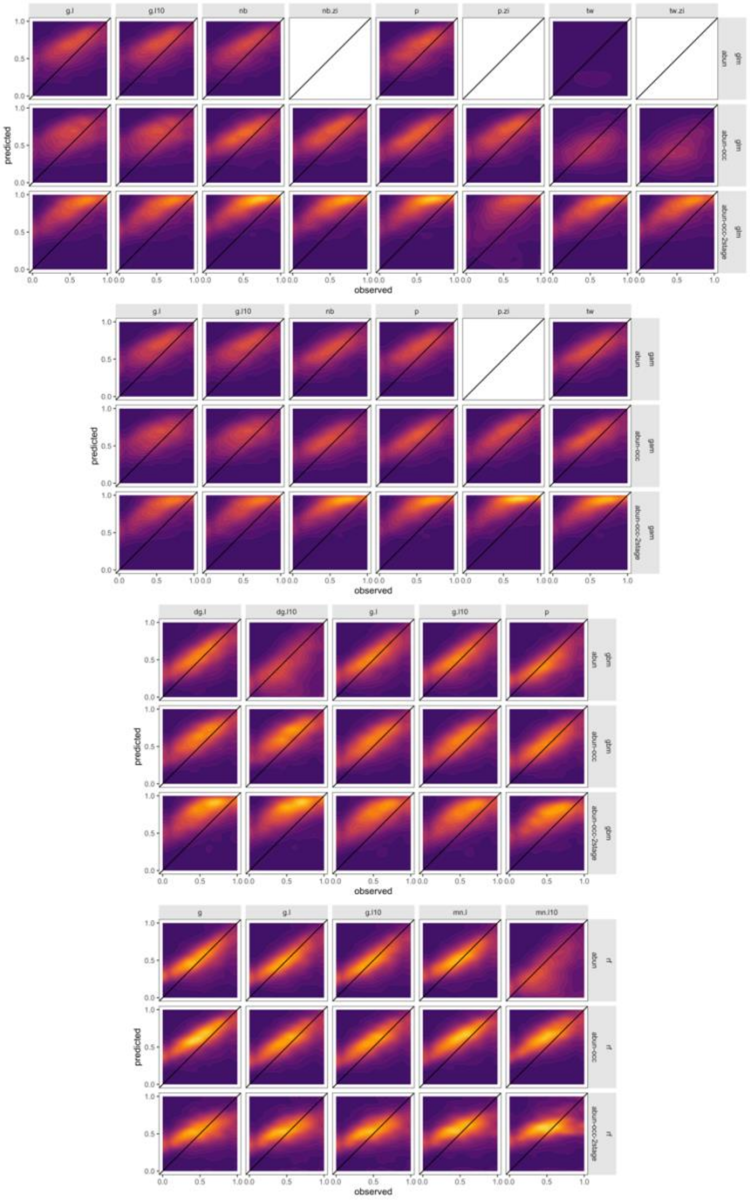

*Figure S15. Comparison between predicted and observed abundance for breeding bird survey within-sample interpolations. Abundance is log10 transformed and rescaled between 0 and 1 to show ability of models to discriminate abundance values. We first removed intercept only models, any predicted values less than 0, and rounded up predicted abundance values less than 1. In addition, we truncate abundance predictions to the lower 1$^{st}$ and upper 99$^{th}$ quantile to avoid extreme values. To avoid common species dominating patterns, for each species, we binned observations into 20 bins and estimated the mean predicted abundance for each observed abundance bin. Note that, due to the log10 transformation, a value of 0 is an abundance of 1. Panel columns represent response data transformations used in models (g = gaussian, g.l = gaussian and log-transformed, g.l10 = gaussian and log10-transformed, dg = first transformed to discrete values but using a gaussian response distribution, nb = negative binomial error, nb.zi = zero-inflated negative binomial error distribution, p = poisson error, p.zi = zero-inflated poisson error, tw = tweedie error, tw.zi = zero-inflated tweedie error, mn.l = multinomial error with log transformation, mn.l10 = multinomial error with log10 transformation).*

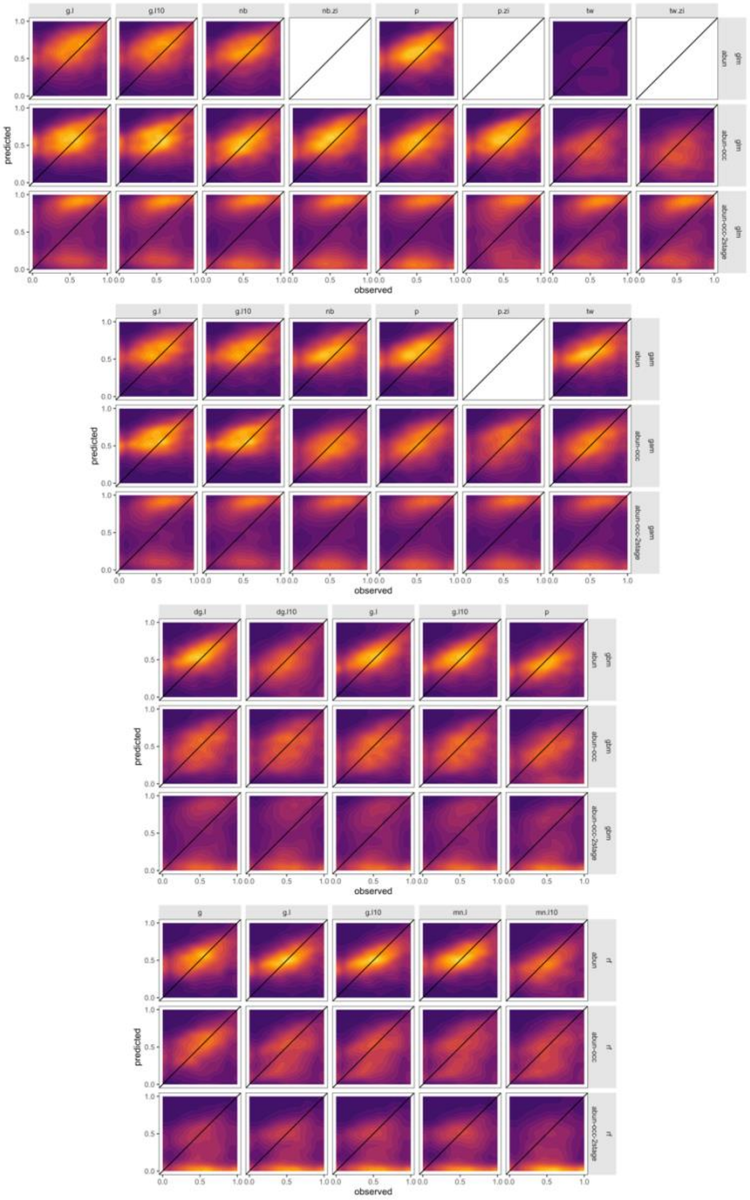


*Figure S16. Comparison between predicted and observed abundance for breeding bird survey out-of-sample extrapolations. For details see Figure S15.*

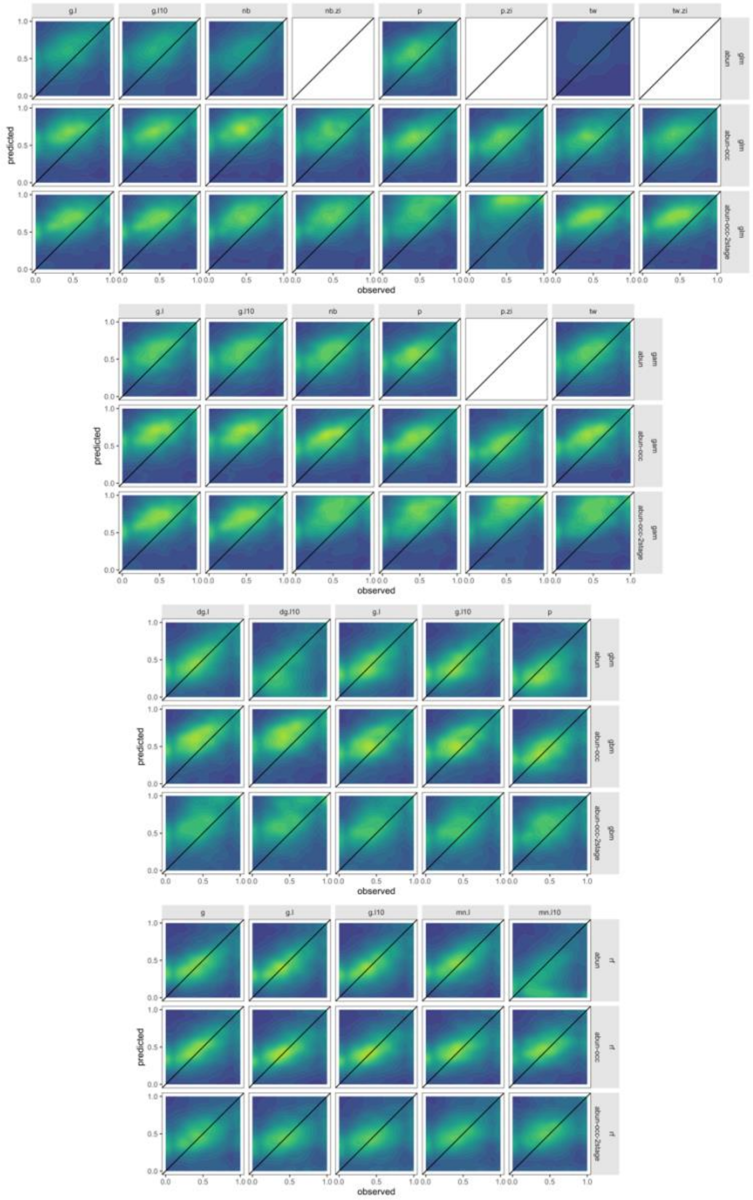


*Figure S17. Comparison between predicted and observed abundance for reef life survey within-sample interpolations. For details see Figure S15.*

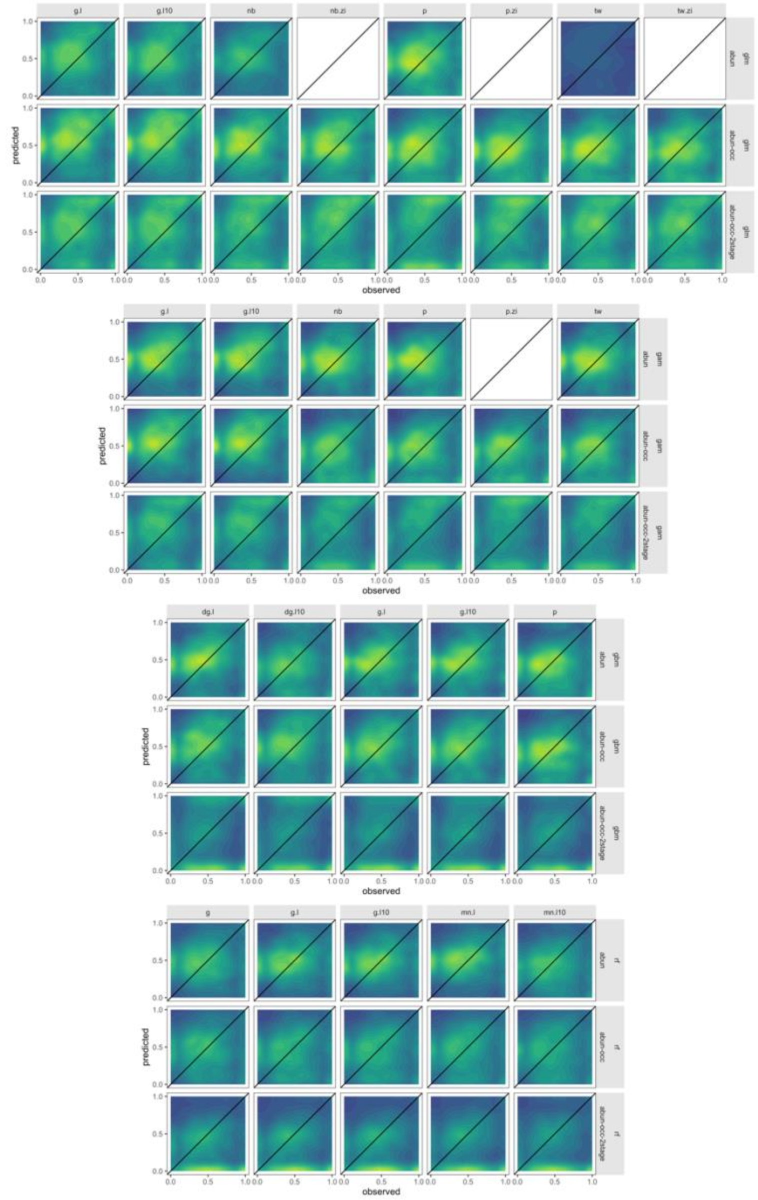

*Figure S18. Comparison between predicted and observed abundance for reef life survey out-of-sample extrapolations. For details see Figure S15.*

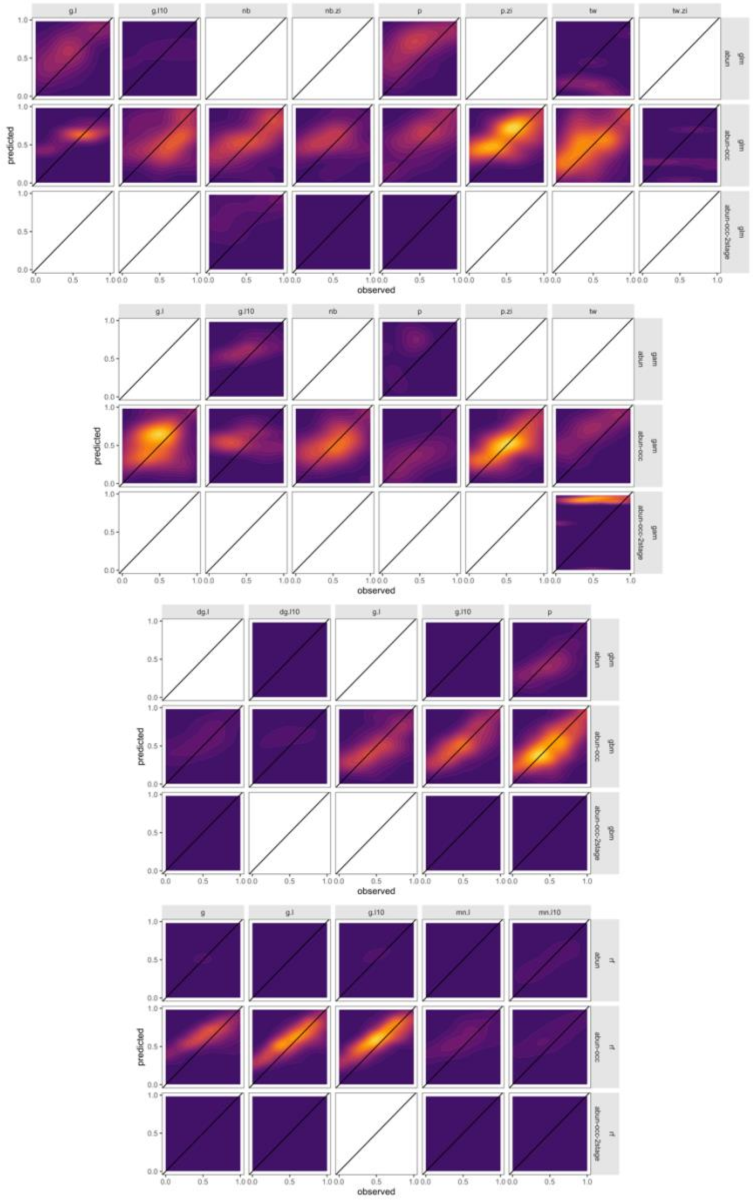


*Figure S19. Comparison between predicted and observed abundance when most discriminatory model for each individual species is selected, for breeding bird survey within-sample interpolations. Most discriminatory model for any given species is variable across modelling frameworks. Model discrimination is determined as the mean model ranks across discrimination metrics. For further details see Figure S15.*

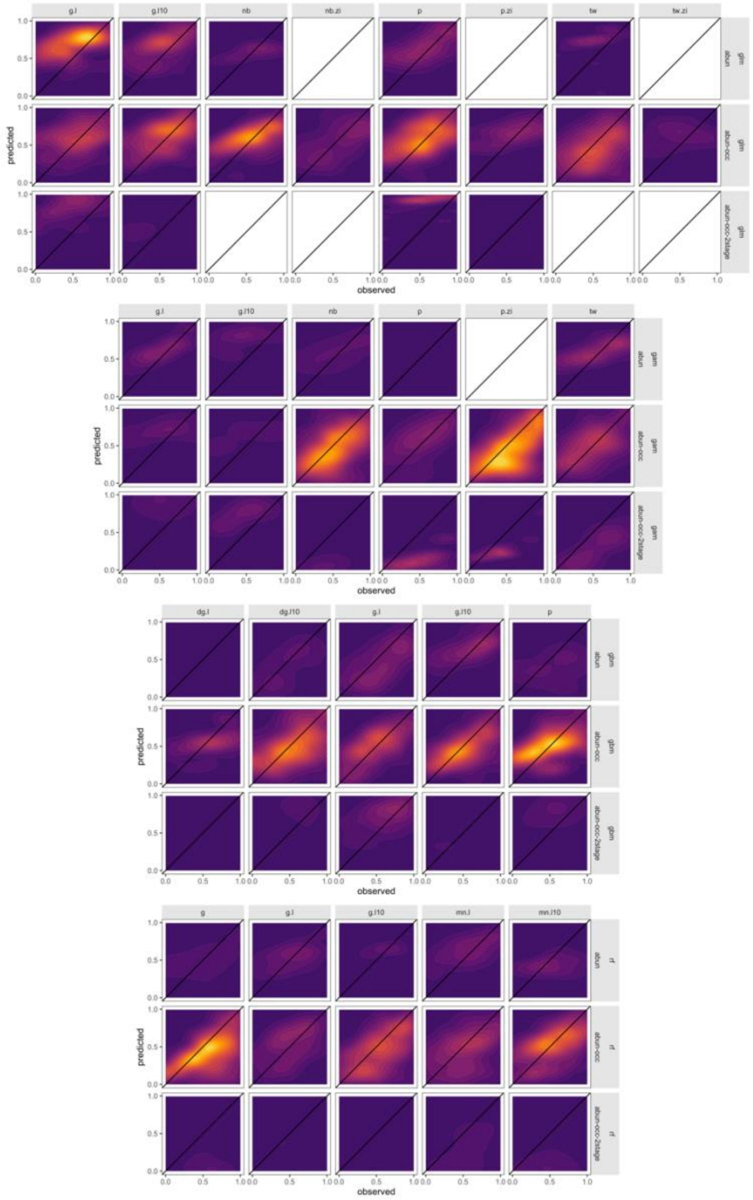


*Figure S20. Comparison between predicted and observed abundance when most discriminatory model for each individual species is selected, for breeding bird survey out-of-sample extrapolations. Most discriminatory model for any given species is variable across modelling frameworks. Model discrimination is determined as the mean model ranks across discrimination metrics. For further details see Figure S15.*

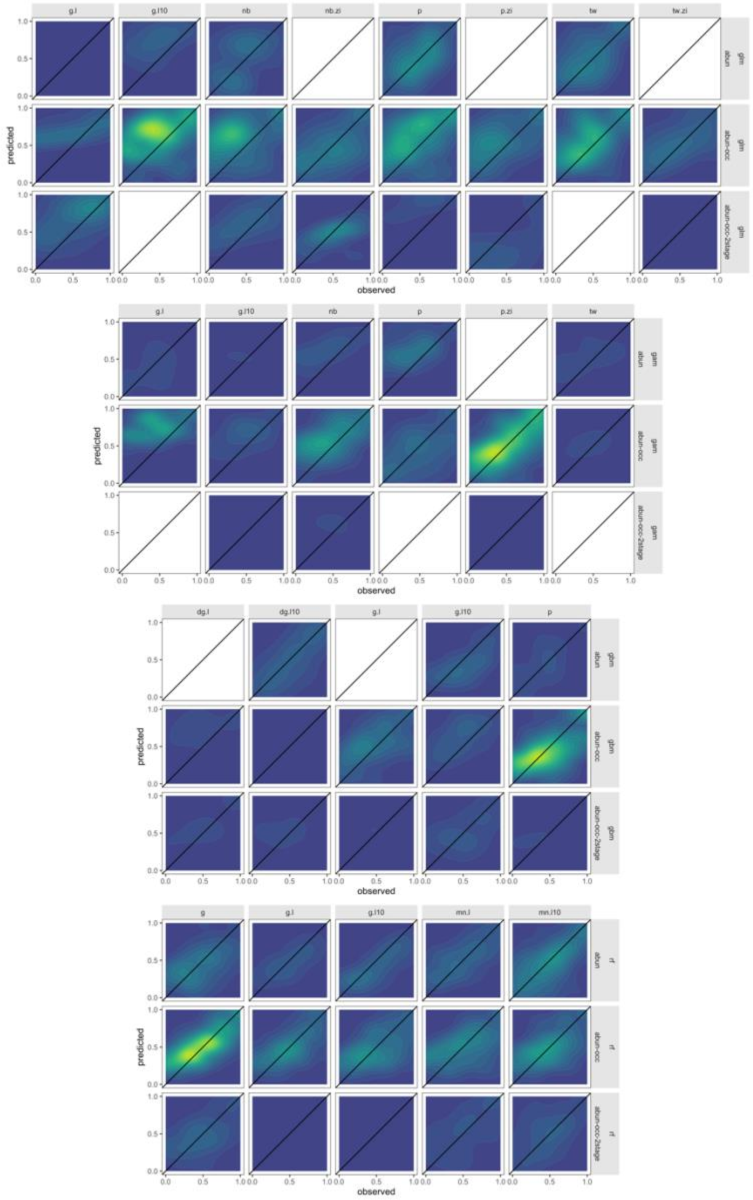

*Figure S21. Comparison between predicted and observed abundance when most discriminatory model for each individual species is selected, for reef life survey within-sample interpolations. Most discriminatory model for any given species is variable across modelling frameworks. Model discrimination is determined as the mean model ranks across discrimination metrics. For further details see Figure S15.*

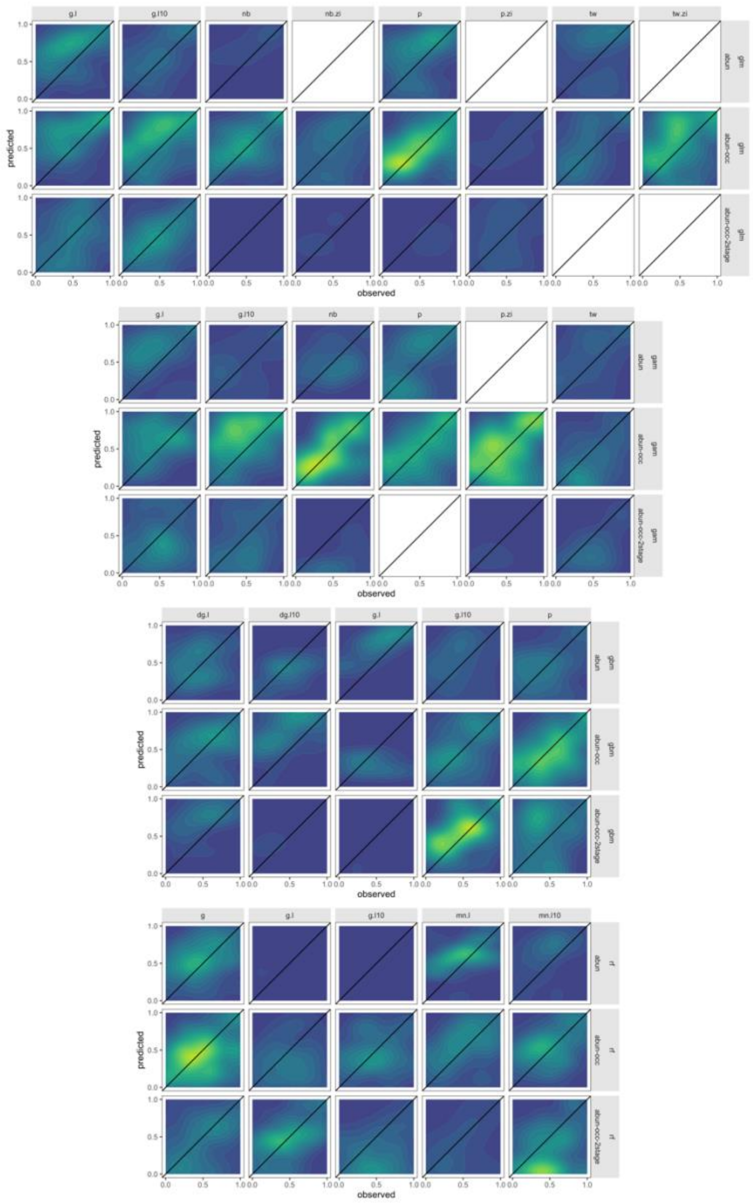


*Figure S22. Comparison between predicted and observed abundance when most discriminatory model for each individual species is selected, for reef life survey out-of-sample extrapolations. Most discriminatory model for any given species is variable across modelling frameworks. Model discrimination is determined as the mean model ranks across discrimination metrics. For further details see Figure S15.*

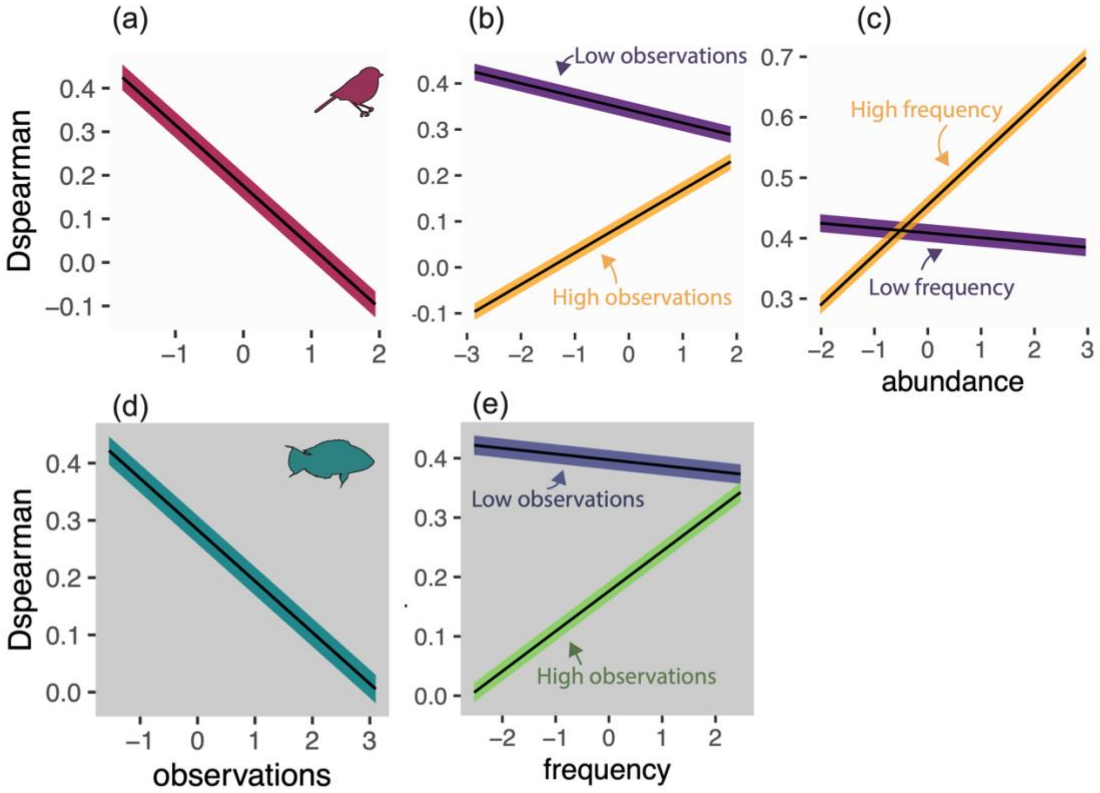


*Figure S23. Effect of species properties on model performance as marginal effects from multiple regressions for out-of-sample model predictions. For full description see Figure 5 in main text.*

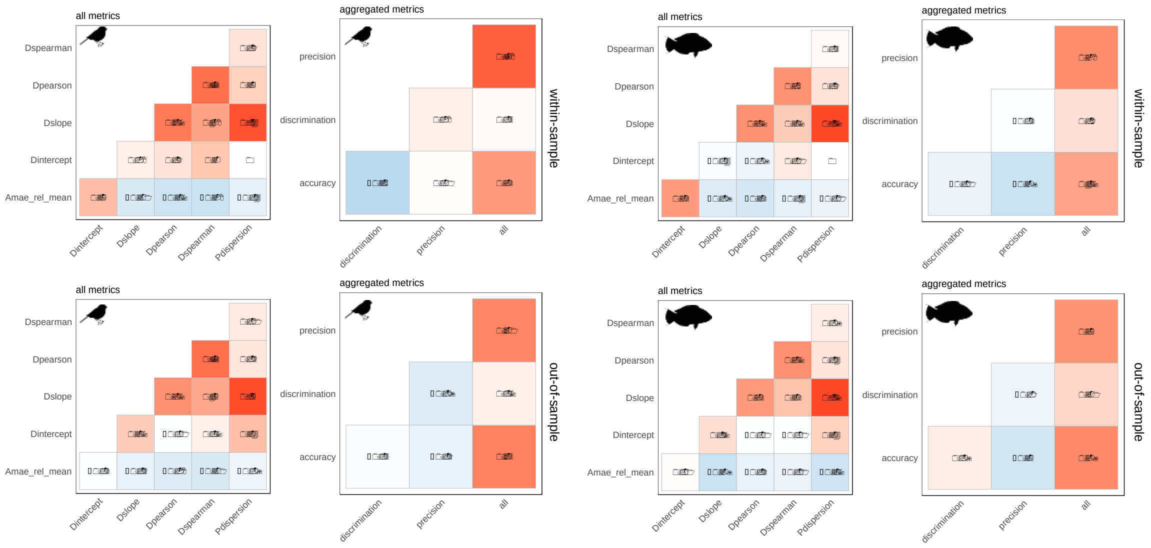


*Figure S24. Correlation amongst performance metrics for within-sample (top row) and out-of-sample (bottom row) cross-validations, across breeding bird and fish datasets and each metric separately and aggregated and ranked metrics.*

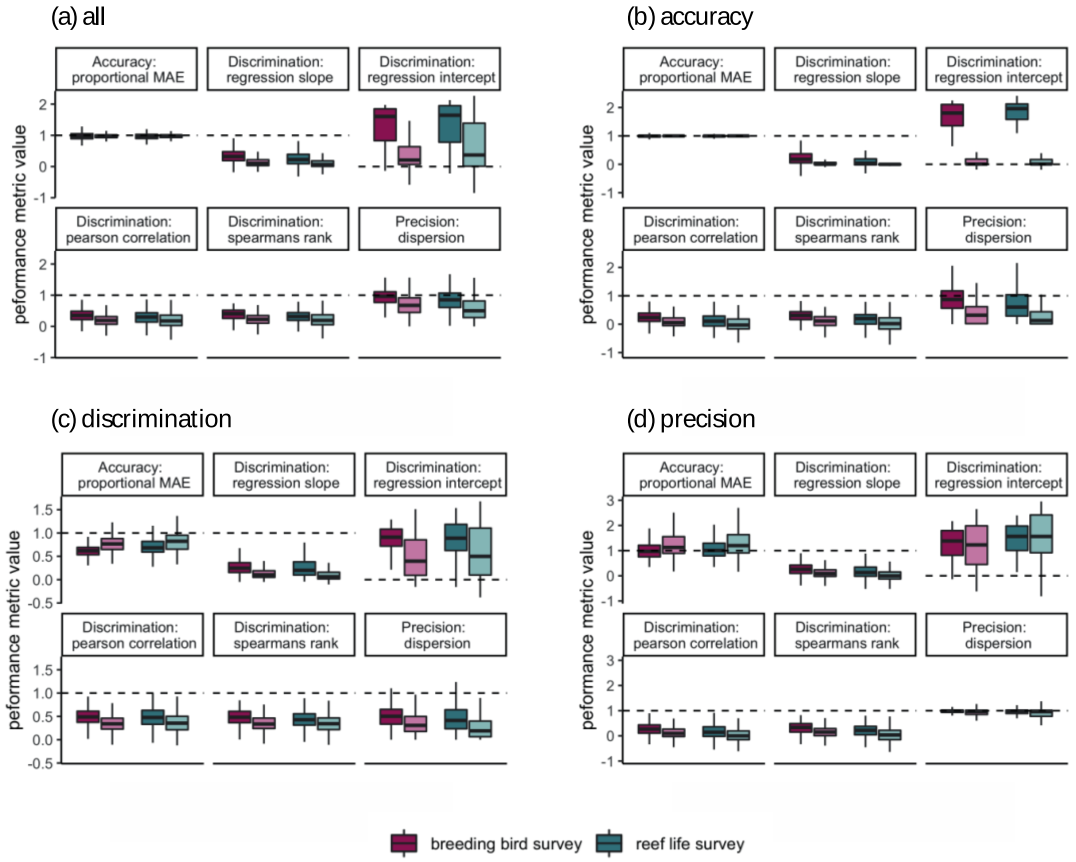


Figure S25. Boxplots of model performance, for each selection of best performing models based on (a) all criteria, (b) accuracy, (c) discrimination, (d) precision for each species across all 6 metrics. Colours indicate breeding bird survey and reef life survey, whereas shading indicates within-sample and out-of-sample projections. Dashed lines indicate target values. Note that the type of is not necessarily the same for a given species in the within-sample and out-of-sample comparisons, as indicated in Figure 2. Central lines correspond to median values, hinges correspond to 25$^{th}$ and 75$^{th}$ quantiles and whiskers correspond to 1.5x the hinges. Outliers are excluded from visualisations.

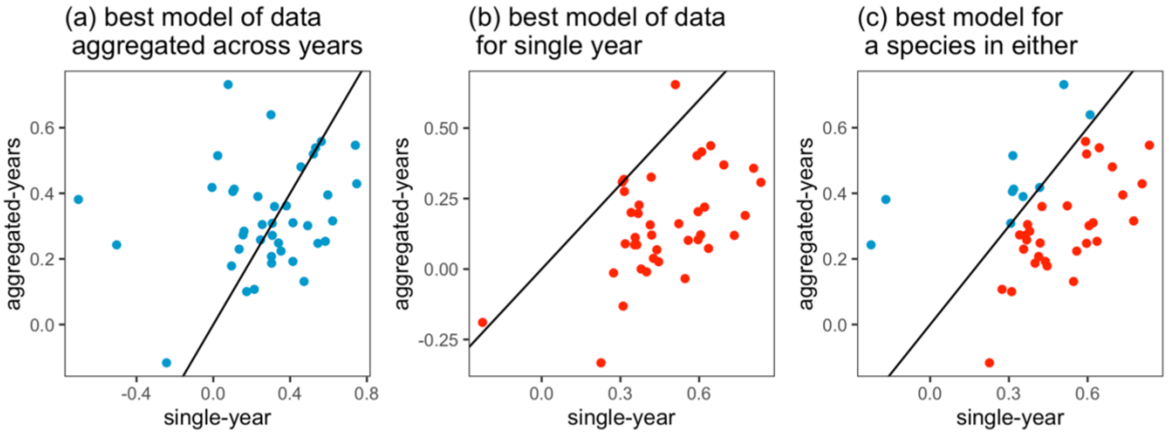


Figure S26. Spearman's rank correlations between observed and predicted abundance per species (points) using 50 species from the breeding bird survey that represent the full range of our data and species characteristics. We compared performance of models fitted using aggregated abundance and climate data from 2007-2017 with performance of models from abundance and climate data from 2017 only. For the climate data we used CRU-TS-4.04 taking the mean, minimum and maximum of: temperature; diurnal temperature range; precipitation rate; vapour pressure; wet days; cloud cover; frost days and potential evapotranspiration, and performing a robust-PCA as in the main manuscript (averaged across 2006-2017 for the 2007-2017 abundance data, and 2017 and 2017-1 for the 2017 abundance data). We compared the best models (in terms of spearman's rank correlation between observations and predictions) for the aggregated abundance-climate data to the yearly abundance-climate data. Points above the solid 1:1 line in a-c represent better models from aggregated abundance-climate data and points below indicate better models from the yearly abundance-climate data. (a) We find that the best aggregated data models had equivalent average performance to the same model fitted to yearly data (mean difference = 0.05, $t = 1.05$, $p>0.05$) and the relationship between the two was weak (rho = 0.19, $p>0.05$). (b) Comparing the best models available using fine-scale data often outperformed equivalent models in the aggregated data models (mean difference = 0.31, $t = 10.18$, $p<0.001$) but performances were significantly correlated (rho = 0.42, <0.01). (c) We found a systematic benefit of using yearly over aggregated abundance-climate data in terms of discrimination (mean difference = 0.12, $t = 3.50$, $p<0.01$). We chose to present the aggregated data in our study because this is the form that most abundance records are available, and as such our results are more generalizable to most ecological studies of abundance-environment relationships. However, we suggest that where available, temporally resolved data on the climatic variability amongst years and its effect on temporally resolved abundances should be modelled. Our interpretation is in general finer temporal data will lead to better models, but some species are still best predicted using more temporally aggregated abundance and climatic data. Both approaches appear valid given the overall spearman's rank values (majority >0.2 in both cases) and future work should address how to improve model predictability where data availability is low. Facets of species ecology that determine the predictability of abundance responses at short-term and long-term time scales should be further explored.
