## Appendix 2 for "A quantitative review of abundance-based species distribution models"

***Appendix 2 – Model fitting details***

*General model fitting procedure*

For all species and models, we balanced the prevalence in our analyses by ensuring that the number of absences was only even a maximum of twice the number of presences in our data. This helps control for differences in the relative number of presences and absences between different species. We randomly sampled absence points using the same random values in R across all species and models. We performed model bootstrapping by refitting models to 10 random subsets of absences to ensure environmental effects from any given random subsample were not spuriously influencing model fits. Note that abundance only models were not bootstrapped as prevalence rate is constant. Model predicted values were produced for our held-out verification and validation sets for each model bootstrap and we quantified the mean, median and standard deviation across bootstrap predictions.

Four transformations of response data were used in total. All predictions were back-transformed before assessment of model predictive ability presented in the main manuscript. Log transformations were applied after the addition of 1 to all values and model predictions were back-transformed by exponentiating *e* to the power of predicted values and taking 1 from these values. For log10 back-transformations were applied by exponentiating 10 to the power of predicted values and taking 1. To transform response data to a discrete scale we first applied log and log10 transformations, next we rounded values upwards and truncated values to a maximum of 6 to avoid having too many classes given the sometimes-low data availability. When discrete data were used, for each bootstrap, we estimated predicted probabilities of each abundance class and estimated the mean predicted value of classes weighted by each class probability. These predicted class values were back transformed as for log and log10.

*Occurrence probability models*

We fitted a set of models with a single covariate of site suitability (two-stage abundance models). We derived site suitability values for each site by first fitting traditional presence-absence species distribution models for each species (i.e., climate envelope, environment niche models). We first converted all abundance values to presences and fitted models to equal sized presence bootstraps such that all locations in our data have a site suitability prediction for each species. We next applied the same routines as for the abundance models (see below) for the same set of statistical frameworks but instead with a binomial response distribution (generalised linear models, generalised additive models, random forests, gradient boosting machines). We ensembled predicted values across all models. We estimated suitability values in verification and validation data without including these data in suitability models to refrain from circularity in using verification and validation data in these suitability models which would inflate predictive power when applied to abundance models.

**Linear and generalized linear models:**

*Conceptual justification*

Fitting complex response curves may result in a model that generalises poorly as models are overfit to environmental and sampling noise which then informs the environment response function (Jones *et al.* 2012; Merow *et al.* 2014; Brun *et al.*

2019). Simpler functions may better approximate the often unimodal shape of species climatic niches, particularly for poorly sampled species (Austin 2007; Boucher-Lalonde *et al.* 2012; Waldock *et al.* 2019).

*Modelling package*
We fitted linear and generalised models in glmmTMB (Brookes et al. 2017; version 0.2.3) using maximum likelihood. glmmTMB was chosen for its flexibility to fit multiple error distributions within one linear modelling framework (See Table S2).

*Details*
We fitted 2$^{nd}$ order polynomial (i.e., quadratic) functions for all covariates to allow simple functions describing the relationship between abundance and environmental conditions. For further simplicity, we did not include interactions between environmental covariates. We performed backward stepwise model selection using likelihood-ratio tests (LRT) and the default settings in the buildglmmTMB function in the 'buildmer' package (Cesko 2019; version 1.2.1). We ensured that covariate linear and 2$^{nd}$ order polynomial terms were blocked together during model selection. Where linear terms, but not 2$^{nd}$ order polynomials, were supported by likelihood ratio tests, we re-fitted model selection including this simplification. Where models included zero-inflation terms, we identified the best covariate structure from the fixed-effect component alone and refitted models with the same model structure in both fixed and zero-inflated model formulas. We applied identical procedures, but not necessarily identical model structures, across all error distributions. We do not filter models by convergence criteria as our primary aim was to fit predictive models for all species (even those with few available data and poor model identifiability) and we infer a lack of convergence as poor parameter identification which will carry through into higher predictive error.

**Generalised additive models:**

*Conceptual justification*
Generalised additive models allow a more complex relationships between the environment and abundance, than quadratic relationships fitted with glms but do not contain as complex interaction structure as machine learning approaches (see below). This flexibility may provide a better trade-off between the non-linear environmental relationships that leading to extreme abundance values and the model fitting to noise in the data (Merow *et al.* 2014).

*Modelling package*
We fitted generalised additive models in using the package 'mgcv' using the function 'gam' which fits models by quadratically penalized likelihood maximisation (Wood 2011, 2017; version 1.8-28).

*Details*
We included all covariates in models and performed model selection using null-space penalization which penalizes the effect of covariates to 0 essentially removing unimportant terms from models (i.e., setting select = T; Marra & Wood 2011). We used thin plate regressions splines and selected a basis dimension (k) of 3 for all covariates to allow intermediate flexibility and computational efficiency (Wood 2017).

We first tested that each covariate alone produces a fitted model and only included covariates in multiple regressions where single covariate models were identifiable.

**Random forests:**

*Conceptual justification*
The drivers of abundance may result from complex and difficult to predict interactions amongst environmental gradients. However, even if the causes of underlying abundances are unknown and complex, they can still be accurately modelled. Random forests model complex interactions amongst predictor variables and often have a high degree of classification accuracy (Cutler *et al.* 2007; Marmion *et al.* 2009). In addition, they are non-parametric, do not make assumptions about the error structure of response data (i.e., non-normality), and are insensitive to outliers which make them suitable for use with abundance data which can show extreme abundance values (highly skewed error distribution).

*Modelling package*
We fitted random forests, a boostrap-based classification and regression tree method, using the function 'randomForest' in the package *randomForest* (Liaw & Wiener; Version 4.6-14).

*Details*
We applied all default settings apart from doubling the number of trees (ntree) to 1000 (Breiman 2001).

**Gradient boosting machines:**

*Conceptual justification*
As in random forests.

*Modelling package*
We fitted gradient boosting machines using the function 'gbm' and the package *gbm* (Greenwell et al. 2019; Version 2.1.5) which use a stochastic gradient boosting strategy (Friedman 2002). The loss function depended on the error distribution modelled in table 1.

*Details*
We first identified the optimal number of iterations in our gradient boosting machines using all observations and predictors in 10-fold cross validation with a bag fraction of 0.8. We chose a high bag fraction, because we have relatively small individual datasets so computation costs of a high bag fraction were not large, in addition a small bag fraction would have increased the chance of a poorly fitting model (Natekin & Knoll, 2013). We applied cross-validation to assess number of trees because it has been shown to be preferable to out-the-bag or independent test set methods available in the same package (Greenwell et al. 2019). We attempted to balance computational feasibility and model adequacy by using an interaction depth of 3, shrinkage (learning rate) of 0.001 and a maximum number of trees (iterations) of 10,000. The optimal number of iterations was assessed using the deviance of observations not used in selecting the next regression tree using the function 'gbm.perf' (note that the exact loss function varies between distributions). 10-fold

cross-validation could not be performed in the package *gbm* with a single covariate (i.e., abundance-2 stage models), in such instances, we identified the optimal number of trees using the function gbm.step in the package *dismo* (Hijmans et al. 2017; version 1.1-4). We used identical parameters as above. We refitted this case of models in *gbm* using the tree number identified in *dismo*. In some instances, models failed to identify the optimal number of trees using 'gbm.step' because the learning rate of 0.001 was too high, as such, we reduced the learning rate by 0.0001 for 500 iterations until an optimal tree number was identified and fitted a final gradient boosting machine using the optimal number of trees described above, a bag fraction of 0.8, an interaction depth of 3 and learning rate 0.001.

**Occupancy Species Distribution Models:**
We used the same four algorithms as above, but for presence-absence responses. For GLM and GAM we used a binomial family distribution, for GBM we used Bernoulli and used RF for classification rather than regression. Variable selection was performed as above. We used the standard settings for RF but set the tree number to 1000. For GBM we set the hyperparameters as follows: n tree to 10,000, interaction depth to 3, shrinkage to 0.001, bag fraction to 0.8 and cv folds to 10. We then used the gbm.perf function to assess the optimal n tree and refitted the model with no cv folds and the n tree optimized. As in the abundance SDMs, we balanced prevalence and bootstrap subsampled absences 10 times and averaged predictions across bootstraps. We produced ensemble predictions of species occupancy across GLM, GAM, GBM and RF as the mean predicted probability of occurrence at a given site.
